# Time-resolved functional rotation crystallography probes protein dynamics during enzymatic catalysis

**DOI:** 10.64898/2026.04.24.718481

**Authors:** J. M. J. Martel, N. Caramello, S. Coquille, E. Mathieu, L. Petit, P. Jacquet, A. Appolaire, F. Leonarski, V. Olieric, M. Wang, D. Madern, A. Royant, S. Engilberge

## Abstract

Kinetic crystallography (KX) encompasses methods designed to characterize structural changes associated with protein dynamics and function. Here, we present Time-Resolved Functional Rotation Crystallography (TR-FRX), an integrated KX workflow combining room-temperature rotation crystallography on single crystals, nanolitre ligand delivery, and optional *in crystallo* optical spectroscopy using conventional synchrotron instrumentation. In TR-FRX, biochemical function is initiated by dispensing a nanolitre volume of ligand onto a single crystal while X-ray diffraction data are collected continuously. The resulting dataset is divided into sequential subdatasets, yielding a series of structural ensembles each averaged over the duration of their data collection time window. Together, these structural ensembles provide an overview of protein dynamics during ligand or substrate binding. We first applied TR-FRX to hen egg-white lysozyme in order to monitor N-acetylglucosamine binding and visualise associated local solvent reorganization. We then applied it to crystals of *Chloroflexus aurantiacus* malate dehydrogenase, first demonstrating by *in crystallo* spectroscopy that they are catalytically active, then revealing by X-ray diffraction ligand-associated solvent reorganization and active-site conformational changes over time windows ranging from hundreds of ms to tens of seconds. TR-FRX provides an accessible single-crystal workflow for establishing conditions in which ligand associated structural changes can be followed at room temperature, thereby generating preliminary data to guide future time-resolved serial crystallography experiments aimed at determining the structure of reaction intermediates.

## Introduction

Enzymes are dynamic molecular machines whose physiological functions rely on transient structural changes (1). Many conformational transitions central to enzymatic catalysis, including ligand binding, loop rearrangements, and domain motions, occur on timescales comparable to enzymatic turnover. These processes frequently unfold within hundreds of milliseconds, although their rates vary across enzyme classes and physiological conditions (2). Capturing such reaction intermediates and associated protein dynamics is therefore essential for understanding enzymatic mechanisms and establishing experimentally supported models of biological function (3). However, conventional structural approaches, including cryogenic crystallography, cryo-electron microscopy and computational structure prediction, predominantly describe stable or highly populated conformational states and often provide limited access to short-lived intermediates (4,5). Kinetic crystallography encompasses a range of complementary approaches for investigating structural changes associated with biochemical reactions and molecular recognition (6,7). Among these, time-resolved macromolecular crystallography (TR-MX) follows structural evolution after a defined trigger using either single-crystal rotation approaches (8,9) or more generally serial data-collection strategies at synchrotrons and X-ray free-electron lasers (10,11,3,12,13). Reactions may be initiated by light excitation, ligand delivery, rapid mixing, temperature changes or chemical perturbations (14). Light-triggered approaches can provide excellent reaction synchronization and temporal resolution extending from femtoseconds to milliseconds for naturally photosensitive systems or crystals prepared with photocaged compounds (3,8,11,9,12,15). Ligand-triggered approaches complement these methods by enabling the study of reactions initiated by diffusible substrates, coenzymes, or effectors. They typically access timescales from hundreds of milliseconds to seconds because their temporal resolution is limited by diffusion through the crystal. In larger crystals, diffusion may generate concentration gradients, heterogeneous occupancies and incomplete synchronization across the X-ray exposed volume (16,17).

Ligand-triggered TR-MX now comprises a diverse range of serial data collection strategies, including mix-and-inject serial crystallography (MISC) (18–21), microfluidic mixing (22), fixed-target methods such as the Liquid Application Method for time-resolved Analyses (LAMA) (23), drop-on-drop delivery (24) and tape-drive approaches (25,26). These methods differ in sample-delivery strategy, crystal requirements, and achievable temporal resolution. By sampling defined reaction delays across many crystals, serial crystallography limits radiation damage while providing high redundancy at each delay, facilitating the detection of weakly populated, short-lived intermediates.

Despite the capabilities of existing time-resolved approaches, establishing suitable experimental conditions remains demanding, particularly when protein functionality, ligand delivery, and the relevant kinetic regime have not yet been characterized *in crystallo*. We therefore developed Time-Resolved Functional Rotation Crystallography (TR-FRX), an accessible single-crystal workflow designed to verify macromolecular function *in crystallo*, identify suitable crystals and crystallization conditions, optimize ligand-delivery conditions and concentrations, and detect structural changes before more challenging serial experiments can be undertaken. TR-FRX combines continuous room-temperature rotation crystallography with nanolitre ligand delivery at a defined delay while diffraction data are collected from a single crystal. Here, “functional crystallography” refers to the determination of successive structures from a crystal in which the macromolecule remains functionally active and the biochemical reaction proceeds *in crystallo*, allowing the resulting structural series to be interpreted in the context of the biological activity (Fig. 1A).

**Figure 1.**
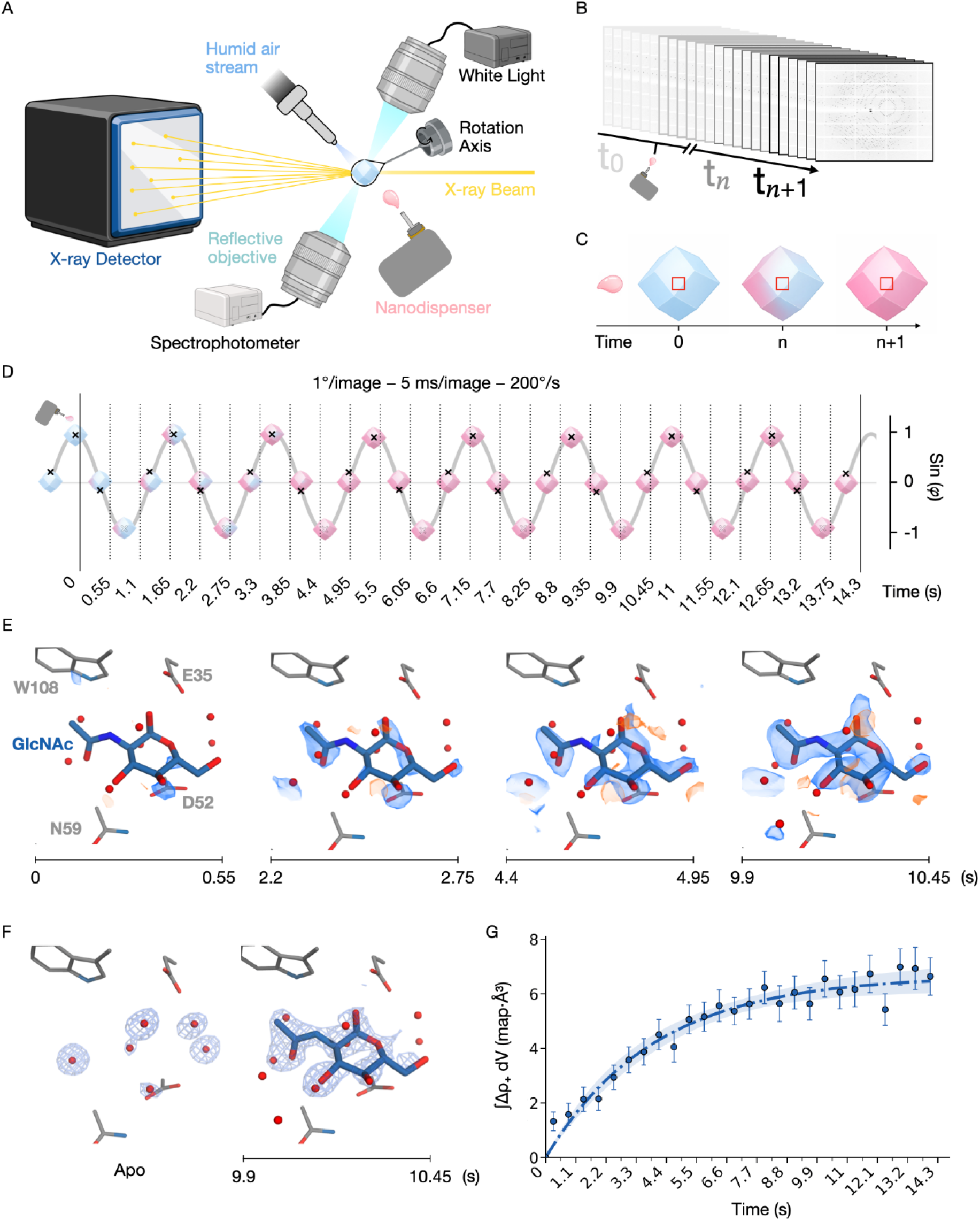
Principle of TR-FRX and proof-of-concept ligand binding. (A) Experimental setup combining in crystallo UV-Vis spectroscopy with X-ray diffraction and precise nanolitre-scale ligand injection onto a single crystal during data collection. (B) TR-FRX continuous data-collection showing a single large rotational data collection acquired continuously and subsequently partitioned into sequential subdatasets, each corresponding to defined time window. (C) Schematic of the crystal region probed by a small X-ray beam (30×30 µm²; red rectangle). A pink droplet denotes substrate delivery, and the pink gradient indicates substrate diffusion in the crystal. (D) Schematic of data collection and subdataset slicing. Trace (sin φ) represents the crystal rotation, where φ = ωt is the crystal rotation angle (1° per image, 5 ms per image, ω = 200 °s^-1^). Crystal icons show successive orientations; the black cross marks a fixed point on the crystal and is dotted when facing away. The left solid vertical line marks injection (t = 0) and the right solid line, the end of data used. Dotted lines mark the boundaries of the 0.55 s subdatasets. (E) F_obs(t)_−F_obs(apo)_ isomorphous difference maps of the active site for subdatasets spanning (0 - 550 ms) to (9.90 - 10.45 s) after GlcNAc injection (contoured at ± 3 σ; positive in blue, negative in orange). (F) Representative 2Fo − Fc electron density maps of the active site before injection (left) and 10.45 s after (right), contoured at 0.8σ. (G) Integrated F_o (t)_ −F_o (Apo)_ difference density within a 1.5 Å mask surrounding the GlcNAc binding site across successive 550 ms subdatasets. The solid curve represents a weighted mono-exponential fit constrained to pass through the origin. The shaded region indicates the 95% confidence interval of the fitted time constant τ = 3.96 [3.15, 5.08] s. See Supporting information for integration and fitting procedure.

TR-FRX can be implemented on standard macromolecular crystallography beamlines by adding a ligand-delivery device to a conventional room-temperature rotation setup. A single droplet of 18 nanolitres is dispensed onto a mounted crystal maintained under controlled humidity during continuous data collection (Fig. 1A). The resulting dataset is subsequently divided into sequential subdatasets, including a pre-injection reference and a series of post-injection subdatasets (Fig. 1B and S1). Each subdataset corresponds to a defined acquisition time window and yields a model representing the diffraction signal averaged over both that interval and the exposed crystal volume (Fig. 1C, 1D). TR-FRX models should therefore be interpreted as time- and volume-averaged ensembles rather than instantaneous or fully synchronized intermediates. Because structural populations may evolve during the collection of a subdataset, too fast or weakly populated species will be missed or underestimated, and refined occupancies are consequently interpreted as apparent, averaged values rather than absolute populations at a given reaction time point (27).

The principal limitation of TR-FRX arises from the timescale of ligand diffusion (17), resulting in asynchronization of the reaction initiation in the whole crystal volume. Additionally, rotational averaging during data collection also limits time resolution, and cumulative irradiation of the same crystal volume needs to be taken into account (28,29). An important advantage, however, is that collecting all subdatasets from the same crystal volume minimizes variability and preserves a high degree of isomorphism throughout the series. The temporal sampling is determined by the duration of each subdataset and therefore depends on crystal symmetry, detector speed, beam size, and photon flux. In contrast, serial crystallography can repeatedly sample defined reaction delays across many crystals and is better suited for resolving faster or weakly populated intermediates. TR-FRX is therefore intended primarily for ligand binding and comparatively slow structural responses in crystals, and as a preliminary workflow for identifying suitable experimental conditions and structural changes before subsequent serial crystallography experiments. Its dense and continuous temporal sampling is particularly well suited to follow slow, large-amplitude conformational rearrangements that involve coordinated movements of multiple protein regions during enzyme function (1,30). In combination with TR-FRX, *in crystallo* UV-Vis absorption spectroscopy provides an independent assessment of crystal activity and helps relate the structural series to the apparent reaction timescale, thereby guiding the choice of crystallographic acquisition windows.

To evaluate potential applications, we selected two benchmark systems. Hen egg-white lysozyme provides a well-established model for monitoring diffusion-driven ligand binding and associated local solvent reorganization (31). We then applied TR-FRX to catalytically active crystals of *Chloroflexus aurantiacus* malate dehydrogenase (*Ca*MDH) (32) to probe protein dynamics associated with enzymatic catalysis. Together, these systems allow us to assess TR-FRX first as a method for following ligand binding then as an approach for investigating slow functional structural changes associated with catalysis over time windows ranging from hundreds of milliseconds to seconds.

### Probing ligand binding dynamics *in crystallo*

As a proof of concept, we assessed whether TR-FRX could monitor diffusion-driven small molecule binding. We monitored N-acetylglucosamine (GlcNAc) binding to HEWL by collecting diffraction data with a 30×30 µm^2^ X-ray beam probing the central region of a large crystal measuring approximately 200×200×200 µm^3^ (Fig. 1C). Four crystals were tested, one of which yielded the complete structural series presented here (Table S1 and S2). Data-quality statistics over the course of the acquisition are presented in Fig. S2. Ligand-associated density was detectable in the first post-injection subdataset, corresponding to a 550 ms time window (Fig. 1E), as revealed by isomorphous difference maps showing the emergence of positive electron density at the canonical sugar-binding site (31,33). No significant difference density features were detected outside of the active site region (Fig. S3). The difference map signal increased progressively across successive subdatasets (Fig. 1E) consistent with a gradual rise in ligand occupancy. Refinement of the initial and late structures (Table S2 and Fig. 1F) further revealed displacement of ordered water molecules from the binding pocket, enabling ligand accommodation and yielding 60% apparent ligand occupancy in the subdataset corresponding to the 9.90-10.45 s time window, in agreement with the expected diffusion timescale for crystals of this size (Table S3).

These results demonstrate that TR-FRX can probe small-molecule binding and associated local structural changes over sub-second to second time windows using a single large crystal at room temperature. The increase in ligand associated difference density was quantified from the structural series (Fig. 1G), yielding an apparent time constant τ of 3.96 [3.15, 5.08] s (see method in supporting information). This value reflects an effective, diffusion-weighted timescale for ligand accumulation in the exposed crystal volume rather than the intrinsic binding rate. This is consistent with probing a large crystal volume, in contrast to cryotrapping studies performed on micrometer-sized crystals where ligand equilibration occurs more rapidly (31).

### Probing ligand-associated structural changes in catalytically active *Ca*MDH crystals on the second to minute timescale

*Ca*MDH catalyzes the reversible interconversion of oxaloacetate (OAA) and L-malate (MLT) with the NADH/NAD⁺ coenzyme pair acting as electron donor and acceptor, respectively. Reduction of OAA to MLT is favored *in vivo* and coupled with NADH oxidation (34). The catalytic turnover rate (k_cat_) of *Ca*MDH in solution is 230 s^-1^ at its optimal temperature of 55 °C, corresponding to ∼14,000 turnovers per minute (4.3 ms per reaction) (32,35,36). This rate is expected to be significantly slower at room temperature and possibly further reduced within the crystalline environment (37). To verify that *Ca*MDH remained catalytically active in the crystalline state, estimate the apparent timescale of the enzymatic reaction under the experimental conditions and optimize ligand and coenzyme concentrations, we first applied *in crystallo* UV-Vis absorption spectroscopy (Fig. 2A and B) on non-rotating *Ca*MDH crystals. The primary purpose of these measurements was to guide the design of the diffraction experiment. We next applied TR-FRX to *Ca*MDH to follow the structural dynamics of the catalytic reaction in active crystals.

**Figure 2.**
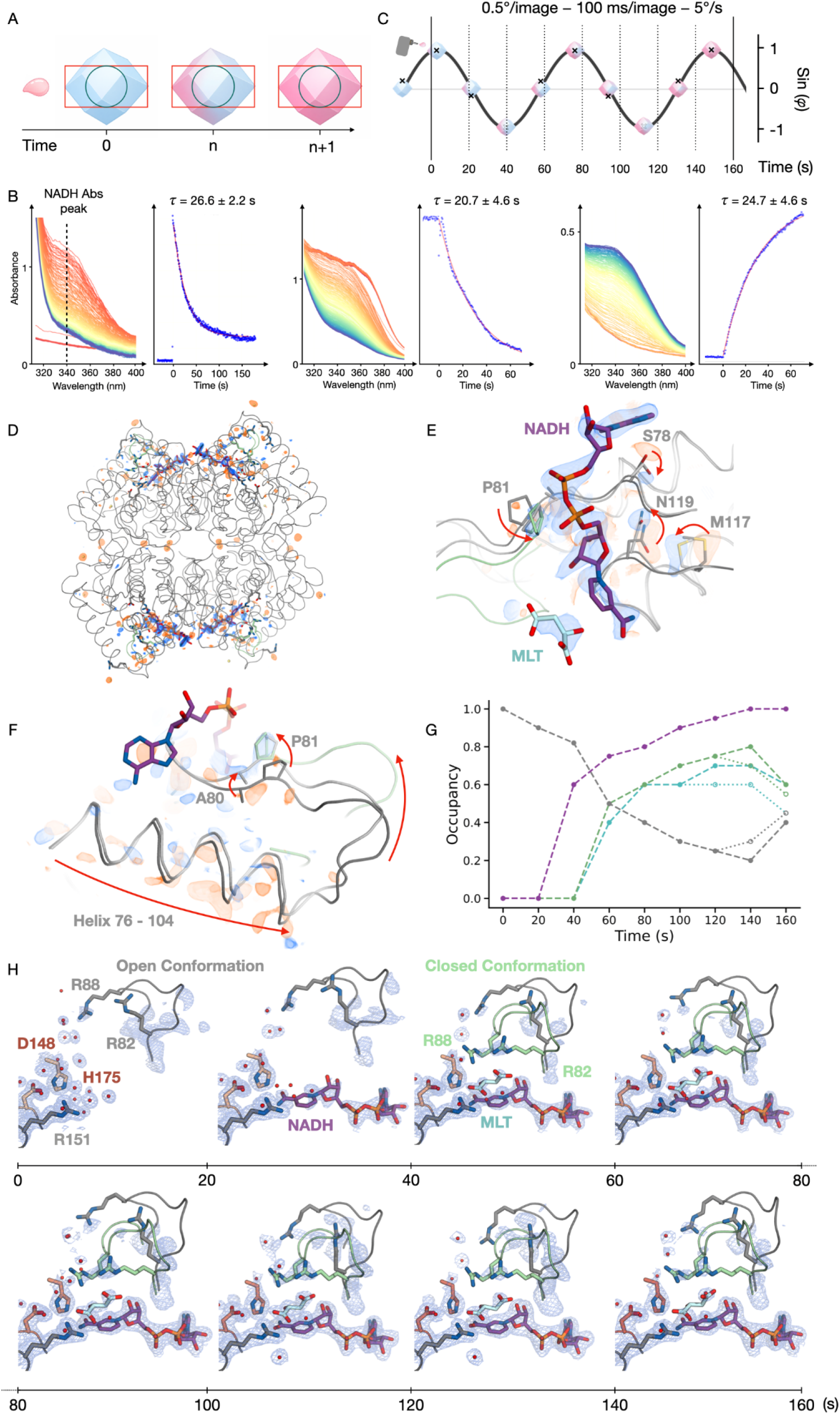
TR-FRX follows CaMDH structural changes during catalysis. (A) Schematic of the crystal region probed by the reference white-light for UV–Vis absorption spectroscopy with a 100 μm diameter spot (blue circle, left) and by a large top hat X-ray beam (200×100 µm²; red rectangle, right). A pink droplet denotes substrate delivery, and the pink gradient indicates substrate diffusion in the crystal (B) in crystallo activity monitored by UV–Vis absorption spectroscopy. (left) Injection of a 50 mM equimolar NADH/OAA mixture. (middle) Crystals pre-soaked with 100 mM NADH, followed by injection of 400 mM OAA. (right) Crystals pre-soaked with 100 mM NAD⁺, followed by injection of 400 mM malate (MLT). Spectral series are color-coded from red (initial) to blue (final). Reaction kinetics were quantified from the time-dependence of the 340 nm absorbance. (C) Schematic of data collection strategy and subdataset partitioning. The trace represents sin φ, where φ = ωt is the crystal rotation angle. Data post-injection were divided into 20 s subdatasets. (D-F) Fo_(100 s)_ − Fo_(apo)_ difference maps showing structural rearrangements associated with ligand/coenzyme binding (±3σ; positive in blue, negative in orange) apo model is depicted in dark grey and holo in light grey, closed loop (holo model) in light green. (G) Evolution of apparent refined occupancies for key catalytic components, including coenzyme (purple), modeled active-site ligand (cyan), and active-site loop extreme conformations (open in grey and closed in green.) (H) Refined active site structures from sequential 20s subdatasets after injection of an equimolar mixture of NADH and OAA (50 mM). Open and closed conformation models of the active-site loop are shown in grey and pale green, respectively. 2Fo–Fc maps are contoured at 0.8σ. Modeled active-site ligand and coenzyme are shown in pale blue and purple, respectively. Key residues are displayed in stick representation.

UV-Vis absorption spectra were recorded continuously before, during and after nanolitre droplet delivery, allowing the progress of the reaction to be monitored over seconds to minutes across the whole crystal volume (Fig. 2B). For these spectroscopic experiments, 13 crystals were used with sizes ranging from 3.5 × 10^-3^ to 2.2 × 10^-2^ mm^3^ (Fig. S4) and were subjected to various soaking strategies (Table S4). Reaction progress was followed from the time evolution of the absorbance at 340 nm. Injection of an equimolar NADH/OAA mixture produced detectable changes in absorbance at ligand and coenzyme concentrations of 50 mM or above. Mono-exponential fitting yielded an apparent time constant τ of 26.6 ± 2.2 s (K_obs_ = 0.0376 ± 0.0031 s^-1^) (Fig. 2B, Table S4 and Supporting Information).

Because ligand diffusion and chemical turnover are convoluted phenomena within the crystal, this value represents a diffusion- and reaction-weighted apparent reaction rate rather than an intrinsic enzymatic constant measured in classical enzymology. Under comparable buffer conditions in solution, the reaction proceeded faster (τ = 12.9 ± 2.8 s, K_obs_ = 0.0775 ± 0.0162 s^-1^), confirming that the overall process is slowed within the crystalline environment (Table S4). Alternative reaction initiation schemes, including coenzyme pre-soaking followed by substrate injection and probing of the reverse reaction direction, yielded similar kinetic behaviour. This similarity between the time constants likely reflects the combined effects of diffusion and catalysis rather than chemical turnover alone (Fig. 2B, Table S4 and Supporting Information).

Across the tested conditions (detailed experimental soaking conditions and analyses of the *in crystallo* UV-Vis absorption data are provided in the Supporting Information and Table S4), apparent *in crystallo* time constants clustered around ∼25 s and showed no clear dependence on crystal size or reaction direction. These measurements established that *Ca*MDH crystals are functional and that the reaction progresses over several tens of seconds. Rotation diffraction data collection is compatible with this timescale, with subdatasets spanning several seconds.

We therefore performed an initial diffraction experiment by dispensing an equimolar mixture of 50 mM NADH and OAA onto a *Ca*MDH crystal 15 s after the beginning of a 350 s data collection (Table S1). A large top-hat X-ray beam (200 x 100 um^2^) was used on beamline BM07 of the ESRF at a flux of 3.7 x 10^10^ photons s^-1^ to probe a crystal volume comparable to that monitored by *in crystallo* UV-Vis absorption spectroscopy while limiting dose accumulation (Fig. 2A) (38). In total, 10 crystals (Table S1) could be measured using the same data acquisition scheme (Fig. 2C, S1C) during a single beamtime shift of 8 hours. The high crystallographic symmetry of *Ca*MDH crystals (*P*3_1_21) allowed the best TR-FRX data collection to be divided into 8 sequential, complete, post-injection subdatasets of 20 s each (Fig. 2C and S1C), with resolutions ranging from 1.7 Å to 2.3 Å (Table S5). Although the total data acquisition extended to 350 s, only the first 160 s post-injection were retained for analysis because of the deterioration in diffraction quality, as assessed from data-quality metrics (Fig. S5). Each subdataset corresponds to an average diffraction weighted dose (ADWD) of ∼7.5 kGy (39) (Fig. S1C).

Structural changes between subdatasets were initially assessed by combining *F*_o_ – *F*_o_ difference maps (Fig. 2D, E and F) together with refinement of each individual structure (Fig. 2G and H). Each structure represents the diffraction signal averaged over a 20 s time window. In the first post-injection subdataset, corresponding to the 0-20 s time window (Fig. 2H), a reorganization of the solvent network was observed, with ordered water molecules in the active-site region progressively vacating their positions. In the 20-40 s time window, the density associated with NADH became clearly visible in the 2*Fo*-*Fc* electron density maps, with an apparent refined occupancy of 60 % for an overall B-factor of 46 Å^2^ (Supporting Information and Fig. S6). Upon NADH binding, ∼15 ordered water molecules (13 in chain A and 18 in chain D) are expelled from the coenzyme-binding cleft (indicated by ≥ 0.4 Å van der Waals overlap between NADH atoms in the 20-40 s time window model and water positions in the apo model). At this stage, the substrate was not yet visible in the catalytic pocket, consistent with a sequential binding mechanism in which the coenzyme binds before substrate (Fig. 2H). In the subsequent 40-60 s time window, density associated with the substrate or product developed in the catalytic pocket and was accompanied by displacement of three additional water molecules which are present in the apo state. Because OAA and MLT differ only by the conversion of a carbonyl group into a hydroxyl group, they are impossible to distinguish in electron density maps at the observed diffraction resolution. The ligand modeled in the active site was therefore assigned according to the direction of the probed reaction, as described in the Methods section, although the ligand-associated electron density likely contains contributions from both species. The ligand-induced solvent rearrangements create a dehydrated pocket promoting tight ligand stabilization. Despite the global reorganization of the active site pocket (Fig. 2E and F), a conserved water molecule persists across all time windows, remaining hydrogen-bonded to the protein backbone and forming an additional hydrogen bond (3 Å) with the substrate or product upon binding (Fig. S7).

Substrate binding was accompanied by redistribution and increased ordering of the catalytic-loop density encompassing residues 81-91. For practical refinement and comparison across the structural series, the active-site loop was represented using two extreme conformations corresponding to the open and closed states. In the pre-injection and first post-injection subdatasets, the loop appears highly dynamic, as indicated by elevated B-factors (Fig. S6), but was nevertheless modeled predominantly in the open conformation. As substrate-associated electron density developed in the active site, the closed conformation became increasingly populated and was characterized by an approximately 7 Å displacement over the active site (Fig. S8). *F*_o_–*F*_o_ difference maps further reveal that residues 79-81, forming the hinge of this loop, move toward NADH and are accompanied by side-chain repositioning (Fig. 2E) together with a broader reorganization of the helix spanning residues 76-104 on the opposite side of the loop (Fig. 2F). The closure of the catalytic loop positions two key residues, Arg82 and Arg88, in proximity to the substrate, where they contribute to its stabilization through direct electrostatic interactions (Fig. 2H). In addition, loop closure creates a solvent-sealed pocket isolated from bulk water (Fig. S9 A-C), thereby promoting catalysis by excluding competing solvent interactions. These observations support an ordering and redistribution of the catalytic-loop ensemble. The apparent refined occupancy of the coenzyme reaches full occupancy in the subdataset 100-120 s, while substrate/product refines to near-full occupancy (∼0.70) at 120-140 s. Notably, the occupancy of the closed catalytic-loop conformation closely tracks that of the bound ligand, consistent with substrate-driven loop closure. In the latest subdataset 140-160 s post-injection, when no more enzymatic activity is observed by *in crystallo* spectroscopy (Fig. 2B), water molecules reoccupy the active site and only low-occupancy ligands can be modeled. Consistently, the closed conformer of the catalytic loop shows increased disorder and a tendency to reopen (opened loop refined at 40% occupancy in the latest time window), consistent with progression toward reaction completion. The coenzyme remains tightly bound at the end of the reaction (Fig. 2H).

Comparison of the two *Ca*MDH monomers present in the asymmetric unit revealed no major difference in ligand occupancy or catalytic loop conformation. The two monomers differ only in the catalytic loop (residues 81–91) of one conformer (chain A), forming an electrostatic interaction with a neighboring symmetry-related molecule. This interaction, involving residues E89 and D90 and their symmetry-related side chains (E89ʹ, D90ʹ), is clearly disrupted during the course of the reaction (Fig. S9 D).

These data show that TR-FRX can follow the progression of ligand-associated solvent and conformational changes in *Ca*MDH crystals over sequential time windows of 20 s on the seconds-to-minutes timescale. The structural series does not resolve individual catalytic intermediates, but still identifies robust changes in active-site hydration, catalytic-loop organization and coenzyme apparent occupancy that happen while the catalysis occurs in the crystalline state.

### Accelerating TR-FRX to a sub-second sampling regime

To extend TR-FRX toward sub-second structural sampling, we tested an accelerated data collection strategy using a smaller X-ray beam (30×30 µm^2^) and a faster detector on beamline X06DA of the Swiss Light Source. The smaller beam, positioned close to the crystal surface (Fig. 3B) reduces the exposed crystal volume and probes a region with a shorter effective ligand-diffusion time, thereby limiting spatial averaging. Because ligand diffusion time in crystals approximately scales with the square of the diffusion distance (16) this strategy is expected to provide earlier access to structural changes associated with ligand diffusion. In parallel, crystals were preloaded with coenzyme so that the reaction could be triggered solely by substrate injection, minimizing delays associated with sequential coenzyme and substrate binding.

**Figure 3.**
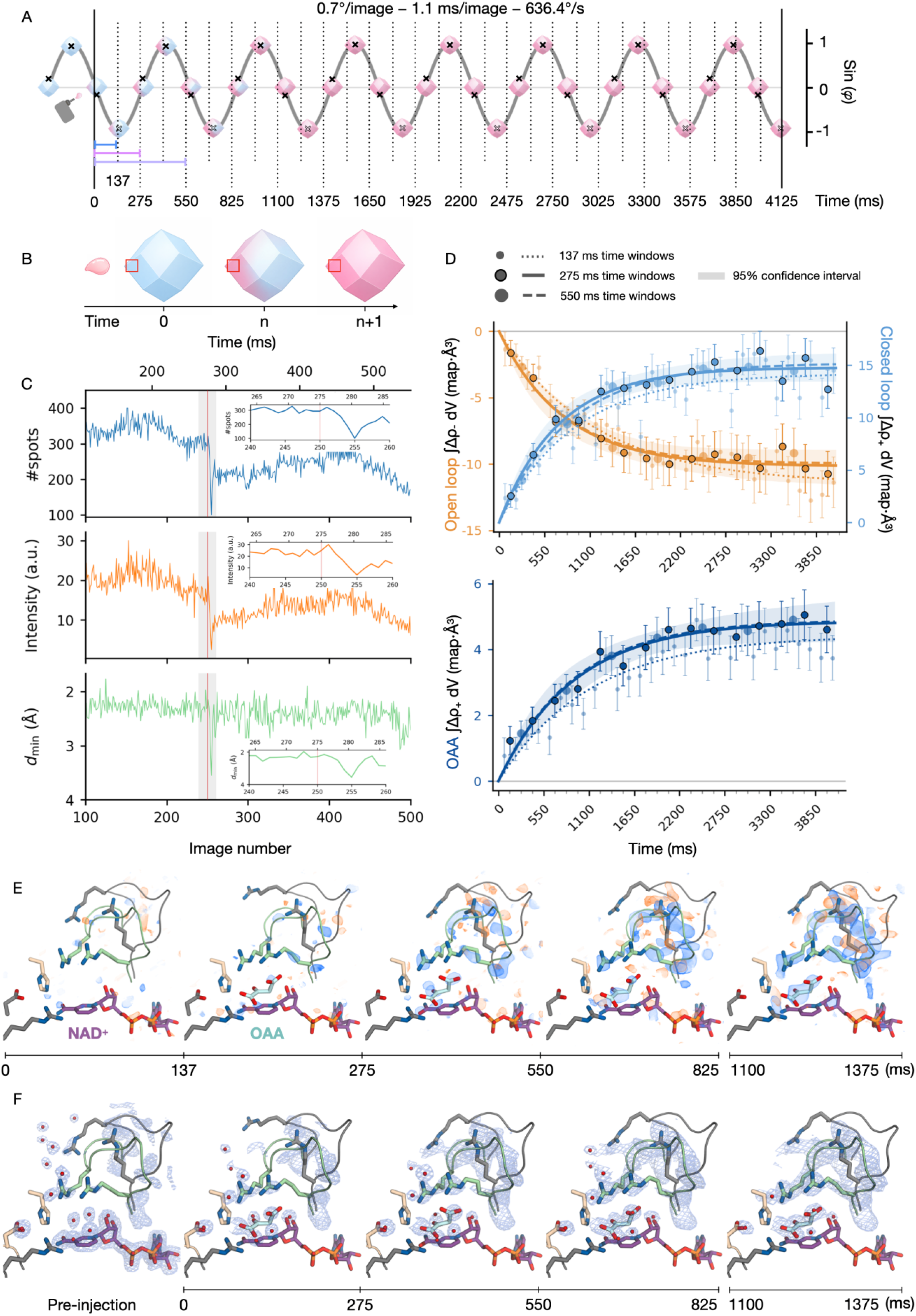
Accelerating TR-FRX to sub-second sampling. (A) Schematic of data collection and subdataset partitioning. The trace represents sin φ, where φ = ωt is the crystal rotation angle. Data were divided into 137, 275 and 550 ms subdatasets (87.5, 175 and 350°), blue, purple and light purple lines respectively. (B) Schematic of the data collection strategy using a small beam (30×30 µm²) covering a restricted region of a crystal. A pink droplet denotes substrate delivery; the pink gradient indicates substrate diffusion in the crystal, the X-ray beam is highlighted by a red square (C) Impact of the injection (red line) on data quality metrics including visible resolution (d min), mean intensity, and number of spots per image. Close up corresponds to the light grey time window around injection. (D) Integrated Fo_(t)_ − Fo_(pre-injection)_ difference density for the open (orange) and closed (light blue) catalytic loop conformations (2 Å radii) and ligand (dark blue, bottom panel, 1.6 Å radius) using 137, 275 and 550 ms subdataset durations. Points show the mean values for chains A and D, with error bars derived from solvent controls. Fits were performed on the two protein chains jointly. The 275 ms series is highlighted with a shaded region indicating the 95% confidence intervals of the monoexponential fits. Corresponding apparent time constants are τ = 0.79 [0.51, 1.18] s, τ = 0.72 [0.53, 1.00] s for open and closed loop respectively and τ = 0.92 [0.59, 1.45] s for substrate. See Supporting information for integration and fitting procedure. (E) Fo_(t)_ − Fo_(pre-injection)_ maps of the active site from 137 to 1375 ms after injection (NCS maps contoured at ± 3σ; positive in blue, negative in orange). (F) 2Fo–Fc electron density maps around the active site (contoured at 1σ).

We monitored the reverse reaction catalyzed by *Ca*MDH (MLT + NAD⁺ ⇌ OAA + NADH). In total, 25 TR-FRX series were obtained from distinct single crystals in 5 hours of beamtime, illustrating the practical throughput of the TR-FRX workflow. The data collection parameters are summarized in Table S1. Data were processed using three alternative subdataset partitioning schemes to evaluate the relationship between temporal sampling and data quality (Fig. 3A and S1D). The finest partition comprised 31 consecutive subdatasets each corresponding to a 137 ms time window and containing 125 images collected over 87.5° except from the reference subdataset consisting in 250 images collected over 175°. The resulting structural series, with a 137 ms time window per subdataset extended from 137 to 4125 ms and yielded spatial resolutions ranging from 1.9 Å to 3.0 Å across the time course (Table S6 and Fig. S11).

Relative to the apo structure (Fig. 2), the NAD⁺-soaked starting state displayed a redistribution of the active-site loop ensemble, with apparent refined occupancies of 65% and 35% for the open and closed limiting conformations, respectively. This suggests that coenzyme soaking alters the conformational equilibrium of the loop before substrate binding. The open and closed models represent here the most extreme conformations selected as a refinement strategy within a broader ensemble, this observation indicates a shift in the loop conformational distribution relative to the apo state (Fig. S8).

Substrate delivery perturbed crystalline order, resulting in a transient drop in diffraction power, and overall diffraction intensity that was maximal ∼5 ms after injection (Fig. 3C). Isomorphous electron difference maps, *F*_o(t)_ – *F*_o(NAD⁺-soaked)_, were computed for each post-injection subdataset spanning from 137 to 4125 ms (Fig. 3E). The strongest reproducible difference density features were located in the active site region (Fig. S12B), with no detectable changes elsewhere in the protein structure. To assess whether these changes could arise from cumulative radiation damage, we performed a control data collection without substrate injection with the same acquisition conditions. Features observed following substrate injection were not reproduced in the control series (Figs. S10 and S12A).

The difference maps revealed positive and negative features around the open and closed conformation of the catalytic loop, together with the progressive appearance of the positive density in the ligand binding pocket (Fig. 3E). These features are consistent with a redistribution of the catalytic loop ensemble toward the closed conformation and with the increasing occupancy of ligand in the active site.

To assess the effect of subdataset duration on map quality, the same diffraction images were also processed as 16 consecutive subdatasets of 275 ms and 7 subdatasets of 550 ms. The principal difference electron density trends were reproduced using all three partitioning schemes, while the 137 ms series showed moderately higher noise, particularly for the weaker features in the substrate/product binding pocket. This comparison indicates that the shortest sampling interval improves temporal discrimination at the cost of lower signal to noise ratio making low-occupancy species more difficult to model reliably. Difference maps were generated using the full data collection (4,000 images) to maximize temporal coverage, whereas structural refinement was restricted to the first 1,500 images, before the onset of substantial deterioration in diffraction quality (Fig. S11).

Integration of the difference density around each loop conformer and around the ligand binding site (Fig. 3D and Fig. S13 and Supplementary Information, 275 ms sampling) gave apparent time constants of 0.79 s (confidence interval: [0.51, 1.18]) and 0.72 [0.53, 1.00] s for the open and closed loop conformers respectively and 0.92 [0.59, 1.45] s for the ligand associated density. Given the magnitude and overlap of the 95% confidence intervals, these values indicate comparable apparent timescales. The 137 ms sampling gave consistent values, whereas the 550 ms series sampled the transition more sparsely, resulting in weaker constraints on the fitted time constant. Structures corresponding to selected time windows (Fig. S11) were refined and apparent occupancies of key catalytic elements were obtained (Fig. 3F). Together, these results demonstrate that accelerated TR-FRX can reproducibly follow ligand-associated active-site and catalytic-loop changes using sub-second time windows.

## Discussion

TR-FRX was developed to enable kinetic crystallography studies on single crystals at room temperature using standard synchrotron instrumentation. In this work, we benchmarked the method on two systems that probe distinct aspects of macromolecular function. Using HEWL, we show that TR-FRX can follow diffusion-driven ligand binding over sub-second to seconds time windows and quantify the associated increase in apparent occupancy from a time series of crystallographic structures. This proof-of-principle experiment establishes that the method is well-suited to follow small-molecule binding and local solvent reorganization in a single crystal, while also providing access to an apparent, diffusion-weighted ligand binding timescale in the crystalline environment. Such measurements may be particularly valuable for room-temperature ligand screening in structure-based drug discovery, where single crystal workflows can reduce sample consumption and simplify experimental implementation (40).

We then extended TR-FRX to the study of enzymatic catalysis using *Ca*MDH, for which the method combines *in crystallo* UV-Vis absorption spectroscopy with X-ray diffraction. This combined approach provided a practical way to estimate the apparent reaction timescale directly in the crystalline state by spectroscopy, optimize soaking conditions, and define acquisition parameters before collecting diffraction data. In the seconds-to-minutes regime, our structural series is consistent with the following sequence of events: coenzyme binding and solvent reorganization precede the appearance of substrate in the catalytic pocket, followed by redistribution of the active-site loop toward a closed conformation and stabilization of the bound ligand (41). The close correspondence between increasing ligand-associated density and loop closure supports a mechanism in which loop motion contributes to ligand stabilization, notably through Arg82 and Arg88. Together, these observations reveal coordinated solvent reorganization and active-site rearrangements during catalysis across successive acquisition time windows, providing a view on the evolution of events associated with protein catalysis within the exposed crystal volume.

The *Ca*MDH system illustrates how several factors, including temperature, ligand diffusion, and crystal packing interactions might influence the apparent reaction rate observed *in crystallo*. The reaction proceeds markedly more slowly in crystals than in solution, likely reflecting the combined effects of decreased solvent dynamics, room-temperature conditions, diffusion constraints within the lattice and stabilizing crystal contacts that can modulate conformational dynamics.

A central methodological point emerging from this study is the importance of matching the acquisition strategy, specifically the probed crystal volume and the subdataset duration, to the kinetic regime of the reaction under study. In TR-FRX, the observed structural evolution reflects a convolution of ligand diffusion with the intrinsic timescales of the enzymatic reaction. As a consequence, different regions of the crystal are not synchronized, leading to a loss of temporal coherence across the crystal and to an effective averaging of structural states that depends on both the exposed volume and the duration of each subdataset. In addition, if populations change during collection of a rotation wedge, different angular ranges can contain different population mixtures. In the initial seconds-to-minutes *Ca*MDH experiments, in which coenzyme and substrate were dispensed simultaneously onto a crystal, the reaction appeared sufficiently slow at the scale of the full crystal volume (τ ≈ 25 s), enabling diffraction measurements with a large beam probing most of the crystal and thereby reporting on the overall structural evolution of the reaction. Under these conditions, the crystal center reaches full ligand saturation after ∼20 s (Table S3), indicating that the earliest post-injection subdataset necessarily contains states resulting from the convolution of diffusion and enzymatic activity, whereas later subdatasets are dominated by enzymatic activity.

For faster events, however, diffusion becomes a strong limitation. Reducing the probed volume with a smaller beam provides a direct route to faster observable time windows. This motivated our accelerated TR-FRX implementation, in which a smaller, brighter beam probes a restricted region near the crystal surface. In this regime, the earliest sub-second subdataset (0 - 137 ms after injection) is acquired before ligand equilibration across the exposed volume, as the characteristic time to reach 50% of the external ligand concentration at the crystal center is ∼180 ms for a 30×30×30 µm³ crystal (Table S3). Consequently, the first post-injection subdataset is particularly affected by ligand diffusion, which generates substantial concentration gradients and heterogeneous ligand occupancy across the exposed crystal volume, thereby attenuating the earliest ligand-associated structural changes. As diffusion proceeds, these concentration gradients are expected to diminish, although each subsequent subdataset still represents a time- and volume-averaged conformational ensemble. In favorable cases, accelerated TR-FRX may nevertheless resolve relatively long-lived transient states, provided that they remain sufficiently populated for a duration comparable to or longer than a single subdataset time window.

A notable experimental consequence of the accelerated TR-FRX measurements is the transient degradation of diffraction metrics immediately after injection (Fig. 3C). This short-lived perturbation is consistent with a mechanical or osmotic shock caused by droplet impact and local solution exchange rather than by a protein conformational transition. This highlights a practical strength of TR-FRX: continuous rotational data collection allows diffraction quality to be monitored frame by frame across the entire acquisition, enabling injection-perturbed frames to be identified and retrospectively excluded during processing while preserving the remainder of the time course. This ability to curate the data, together with continuous sampling of the reaction timeline, facilitates both optimization of subdataset duration and downstream analysis of structural evolution.

We also observe an influence of crystal packing on catalytic-loop dynamics through a Cd^2+^-mediated crystal contact that is abolished in one chain during turnover (Fig. S9 D,E). Together with ligand diffusion effects, these results emphasize that TR-FRX monitors protein dynamics as it unfolds within the crystalline environment, where the observed signal reflects a convolution of crystal packing effects, diffusion, radiation damage and reaction kinetics rather than purely intrinsic enzymatic dynamics.

Radiation damage must be carefully considered during room-temperature diffraction experiments. (29,42–44) Serial synchrotron approaches typically mitigate this by distributing the absorbed dose over many microcrystals, whereas rotation-based methods progressively concentrate the dose within a single crystal. As a rotation-based method, TR-FRX therefore needs to be implemented under low-dose conditions to limit X-ray induced perturbations of catalysis while preserving high-resolution diffraction throughout the time course. In all of our experiments on *Ca*MDH, only ∼4 to 16 kGy (Table S1 and Fig. S1B,C,D) were absorbed per subdataset, resulting in total accumulated doses for the entire refined time series below the ∼140 kGy generally considered acceptable for complete room-temperature datasets. (29,43) This limited dose budget reduces, but does not eliminate, the risk of specific radiation damage and beam-heating artefacts while enabling multiple sequential subdatasets to be recorded from the same crystal. Isomorphous difference maps from dose-matched control data collection without injection were clearly distinct from those obtained after ligand injection and did not reproduce the active-site solvent rearrangements or catalytic-loop changes observed in the TR-FRX series (Fig. S12). The control series nevertheless showed weak negative difference density at several Cd^2+^ sites on *Ca*MDH surface. An additional progressive reorientation of His175 was observed during the slow-series acquisition. Because cumulative irradiation may contribute to this change, it is reported as a structural observation (Fig. S15) but is not used to support the mechanistic interpretation. Practically, TR-FRX complements serial approaches, by combining dose saving with continuous quality control on a small number of crystals.

The present implementation calls for further improvement. Sub-100 ms time windows may become accessible by combining smaller crystals, smaller beam sizes, higher flux densities, and faster detectors. Alternatively, merging incomplete subdatasets from a handful of crystals may provide a practical route to higher temporal sampling.

In summary, TR-FRX fills a methodological gap between cryo-trapping and serial time-resolved crystallography. It retains the accessibility and low sample requirements of conventional room-temperature rotational data collection while introducing time resolution, continuous sampling, and direct compatibility with *in crystallo* spectroscopy. By enabling ligand binding and catalytic structural changes to be followed on single crystals at room temperature, TR-FRX provides an accessible route to ‘functional crystallography’ in which successive protein structures are interpreted within the dynamic context of the measured biological activity.

## Materials and Methods

### Sample preparation and crystallisation

HEWL was purchased from Roche (catalog no. 10837059001) and crystallized at 20 mg mL^-1^ by hanging-drop vapor diffusion against a reservoir solution containing 100 mM sodium acetate (pH 4.3) and 0.8 to 1.3 M NaCl. Crystals with typical dimensions of approximately 200 × 200 × 200 µm^3^ grew within 2 days and belonged to space group *P*4_3_2_1_2.

Overexpression of *Ca*MDH was carried out following similar protocols already described by Dalhus *et al.* (2002) and Talon *et al.* (2014).(36,45) Bacterial cells were lysed by sonication in 50 mM Tris-HCl (pH 7.0), 50 mM NaCl (Buffer A). Supernatant was acidified to pH 6.0 using Bis-Tris propane pH 6.0 before a heat shock at 70 °C for 30 min. Unfolded *E. coli* proteins were removed by centrifugation at 39,000 g for 20 min. The remaining supernatant was dialyzed against Buffer A overnight and loaded onto a Q-sepharose column. *Ca*MDH was eluted with a linear gradient from 50 mM to 1 M NaCl. Fractions containing the enzyme were pooled and supplemented with (NH₄)₂SO₄ for a final concentration of 2 M, then loaded onto a Butyl Sepharose column. Elution was performed using a negative linear gradient, from 2 to 0 M of (NH_4_)_2_SO_4_ in Buffer A. Active fractions were pooled, concentrated, and loaded onto an Enrich 650 size exclusion chromatography column (Bio-Rad), and eluted with Buffer A. Resulting fractions were concentrated to 20 mg.mL^-1^ and stored at 4 °C. Cubic-shaped trigonal (*P*3₁21) crystals (30 x 30 x 30 to 200 x 200 x 100 µm^3^) containing 52 % solvent were obtained in 48 h at 20 °C by hanging-drop vapor diffusion using a mother liquor containing 20 mM CdCl₂, 12 to 24 % PEG400, and 100 mM sodium acetate at pH 4.6 or HEPES pH 7.0 (Table S1).

### TR-FRX experimental setup

TR-FRX experiments were conducted at room temperature (20 °C) on the BM07 beamline at the European Synchrotron Radiation Facility (ESRF) and on the X06DA (PXIII) beamline at Swiss Light Source 2.0 (SLS2.0).(46,47,38) Single crystals were directly mounted in nylon loops (Hampton Research) on the beamline diffractometer in the experimental hutch. To prevent desiccation, crystals were maintained at a controlled relative humidity of 98 %, determined experimentally, using an HC-Lab humidifier (Arinax). Single droplets of 18 nL of ligand solutions (Table S1) were delivered in a non-contact manner directly onto the mounted crystals using a PIPEJET® nanodispenser (Biofluidix/Hamilton) installed in the sample environment and mounted on manual X, Y, and Z translation stages. Alignment of the 500XL injection pipe from the nanodispenser was achieved using the on-axis beamline microscope. This setup (Fig. S14) is inspired by the Acoustic Levitation Diffractometer on the X06SA beamline at the Swiss Light Source (Villigen, Switzerland).(48,49) A single droplet was applied to each crystal at a defined delay during a single continuous rotation data collection, the resulting images were subsequently partitioned into sequential subdatasets.

### Solution and *in crystallo* UV-Vis absorption spectroscopy, data collection and processing

UV-Vis absorption spectra were recorded on beamline BM07 using the online microspectrophotometer from the *ic*OS lab (Fig. S14A), mounted in the sample environment.(46,50,38,51) The incident white light was provided by a DH2000-BAL lamp (Ocean Optics) connected to the setup via a 400 µm optical fibre and focused at the sample position with a 4x reflective objective. The transmitted light was collected via the second reflective objective and transmitted via a 600 µm fibre to a QEPro spectrophotometer (Ocean Optics). Optimal crystal orientation was achieved by rotating the crystal on the goniometer axis until the signal-to-noise ratio of the spectrum at 280 nm was maximal. Spectra were recorded continuously at 500 ms intervals. The injection trigger defines t_0_ and corresponds to manual dispensing of a single 18 nL ligand droplet through the Biofluidix graphical user interface controlling the PIPEJET® nanodispenser. Absorption spectra were processed using the *ic*OS toolbox (52). Smoothing and constant baseline correction (600 - 700 nm) were systematically applied to the series of UV-Vis spectra. Time constants (τ, 1/e) for the enzymatic reaction were determined by fitting monoexponential decay models to the absorbance data at 340 nm. Details of the spectroscopic data analysis are provided in the Supporting Information.

### X-ray data collection and processing

Diffraction images were collected on the BM07 beamline at the ESRF using a Dectris PILATUS2 6M detector.(46) The TTL signal from the detector was duplicated and routed through a BNC505 delay generator (Berkeley Nucleonics Corporation), which was connected to the PIPEJET® nanodispenser controller. Ligand injection via the PIPEJET® nanodispenser was triggered after a controlled delay following the start of data collection, with the precise timing set using the delay generator (Fig. S1A). High-rate data acquisition was performed on the X06DA beamline (SLS 2.0) using a Dectris PILATUS4 2M detector.(47) Timing was controlled with a PandABox to synchronize the detector, shutter, and PIPEJET® nanodispenser, using the position-synchronized output (PSO) of Aerotech A3200 goniometer from the rotation axis as the master signal. Data collection parameters are reported in Table S1 below. During each TR-FRX series, successive subdatasets were collected from the same exposed crystal volume as the crystal underwent repeated rotations without any translation. Table S1 and Figure S1B,D report the data collection parameters together with total exposure time, number of rotations and absorbed dose per subdataset. A control data set without injection was collected on *Ca*MDH at X06DA under the same data collection strategy and therefore dose rate to assess radiation damage effect independently of ligand delivery (Fig. S10 and S12).

Each data collection was divided into sequential complete subdatasets using autoPROC.(53) The first subdataset, corresponding to the structure before ligand injection, was processed independently and used as the reference for subsequent subdatasets. Dedicated scripts used to automate the partitioning of the continuous data collection into sequential subdatasets are available on GitHub (https://github.com/jmartelX/TR-FRX-autoprocessing). Structures were solved by molecular replacement using DIMPLE in CCP4,(54,55) followed by manual model building with COOT(56) and refinement with PHENIX.REFINE.(57) For *Ca*MDH, the catalytic-loop ensemble was represented using two extreme models corresponding to open and closed conformations. Their apparent occupancies, together with those of the coenzyme and active-site ligand, were refined using a consistent protocol while monitoring B-factor consistency across successive subdatasets. These apparent occupancies are used to compare relative trends and should not be interpreted as absolute populations at discrete times. Because OAA and MLT cannot be distinguished unambiguously, the active-site ligand was assigned according to the reaction direction, MLT for the forward reaction and OAA for the reverse while acknowledging that the densities contain contributions from both species. Isomorphous difference maps were generated using PHENIX (see Supporting Information for details). Final models were validated using MolProbity before deposition in the Protein Data Bank. (58)

## Data availability

Raw diffraction images have been deposited in Zenodo under DOI https://doi.org/10.5281/zenodo.19223424 and https://doi.org/10.5281/zenodo.19232847. All atomic models have been validated and deposited in the Protein Data Bank, corresponding accession codes are provided in Tables S2, S5 and S6.

## Author contributions

The initial concept of the method was developed by S.E., V.O. and M.W. *Ca*MDH purification and crystallization were performed by J.M.J.M., S.C. and D.M. The TR-FRX setup was implemented on the BM07 and X06DA beamlines by S.E., P.J., L.P., E.R.M., A.A., F.L. and V.O. S.E. and J.M.J.M. performed data collection, data processing, model building and structure refinement. N.C. provided support for data processing. S.E. and J.M.J.M. validated and analyzed the structures. J.M.J.M. and S.E. prepared the figures. S.E. wrote the manuscript with input from A.R., N.C., D.M., V.O., M.W. and J.M.J.M. All authors approved the final version of the manuscript.

## Acknowledgements

We acknowledge the French Biology/Health Panel Review Committee for provision of synchrotron radiation beamtime at the ESRF (Grenoble, France) on beamline BM07, supported by the French ANR PIA3 (France 2030) EquipEx+ project MAGNIFIX under grant agreement ANR-21-ESRE-0011. This work used the *ic*OS platforms of the Grenoble Instruct-ERIC center (ISBG; UAR 3518 CNRS–CEA–UGA–EMBL) within the Grenoble Partnership for Structural Biology (PSB), supported by FRISBI (ANR-10-INBS-0005-02) and GRAL, financed within the University Grenoble Alpes graduate school (Ecoles Universitaires de Recherche) CBH-EUR-GS (ANR-17-EURE-0003). This work was supported by the Agence Nationale de la Recherche (ANR) under the JCJC program project Catal-X ANR-25-CE11-5075 and AlloSpace ANR-21-CE44-0034-01. The authors acknowledge the Paul Scherrer Institute for beamtime provision under proposal 20250795 and the staff of the X06DA beamline at SLS for support. Sylvain Engilberge thanks his former colleague at the Swiss Light Source Takashi Tomizaki for the initial discussions on the use of the PipeJet nanodispenser for *in situ* mixing of ligands. Tristan Wagner is acknowledged for discussion and advice. We gratefully acknowledge Manuel Maestre Reyna, Marco Bellinzoni, Ahmed Haouz, Claude Didierjean, and Mathieu Schwartz, the first users of TR-FRX on the BM07 beamline whose early experiments on different targets and feedback helped refine the method. Julien Martel acknowledges the CEA for funding his PhD thesis through the CFR program.

## Supporting information

### *in crystallo* optical spectroscopy of the *Ca*MDH system

For a typical *in crystallo* UV-Vis absorption spectroscopy experiment, a single *Ca*MDH crystal was manually mounted in a nylon loop (Hampton Research) and positioned on the goniometer of the BM07 beamline. The crystal was centered within the focal spot of the UV-Vis microspectrophotometer and oriented to maximize the protein absorbance signal at 280 nm while minimizing the baseline absorbance. Absorption spectra were recorded continuously with a temporal sampling of 500 ms with no readout time since spectra acquisition is buffered. After sufficient pre-injection spectra had been accumulated, a single 18 nL droplet of an equimolar NADH/OAA mixture at concentrations ranging from 5-100 mM was dispensed using the Biofluidix graphical user interface controlling the nanodispenser. All spectroscopic measurements and associated experimental conditions are summarized in Table S4.

No detectable catalytic activity was observed below 20 mM ligand concentration, whereas clear kinetic traces were obtained at 50 mM or above. Reaction-associated absorbance changes were fitted with a monoexponential function, from which the characteristic time constant τ and the corresponding apparent observed rate constant K_obs_ were determined. These measurements yielded a characteristic time constant τ = 26.6 ± 2.2 s, corresponding to K_obs_ = 0.0376 ± 0.0031 s^-1^ across crystals with volumes ranging from 3.5 x 10^-3^ to 2.2 x 10^-2^ mm^3^. Because ligand diffusion and chemical turnover occur simultaneously within the crystal K_obs_ is interpreted as an apparent rate constant describing the overall spectroscopic evolution rather than an intrinsic catalytic rate constant.

Additional measurements were performed under comparable buffer conditions in solution using the purified enzyme (20 mg.mL^-1^) mixed with the crystallisation solution to mimic crystalline chemical environment. For these measurements, instead of mounting a crystal in the nylon loop, solution of the enzyme was placed in the loop and subjected to the same acquisition protocol described above, with absorption spectra recorded continuously at 500 ms intervals before, during and after ligand injection. The reaction proceeded more rapidly (τ = 12.9 ± 2.7 s, K_obs_ = 0.0775 ± 0.0162 s^-1^) than in the crystalline state. The slower apparent reaction rate observed *in crystallo* compared to the one measured in solution likely reflects, at least in part, diffusion limitations within the crystalline environment.

Alternative reaction-initiation strategies were also tested by manually pre-soaking crystals with 100 mM NADH before substrate injection with nanodispenser, allowing coenzyme binding to precede reaction initiation. Subsequent injection of OAA at concentrations between 100 and 400 mM produced similar reaction kinetics. For example, injection of 200 mM OAA yielded τ = 31.6 ± 3.9 s, K_obs_ = 0.0316 ± 0.0039 s^-1^ while increasing the OAA concentration to 400 mM accelerated the reaction slightly (τ = 20.7 ± 4.6 s, K_obs_ = 0.0482 ± 0.0109 s^-1^). The reverse reaction was also investigated by pre-soaking crystals with 100 mM NAD⁺ followed by injection of 400 mM malate, yielding τ = 24.7 ± 4.6 s, K_obs_ = 0.0405 ± 0.0074 s^-1^. Overall, reaction kinetics were remarkably consistent across ligand-soaking strategies, crystal sizes, and reaction directions.

### Radiation damage induced structural changes?

During refinement of the slow TR-FRX series, we observed a progressive motion of His175 across the structural series. We therefore analyzed the nine refined structures spanning 0 - 160 s, corresponding to 20 s sampling intervals (Fig. S15). The orientation of the His175 imidazole ring was quantified using the atoms ND1, CG, CE1, NE2, and CD2 to define the ring plane. A normal vector to this plane was calculated for each structure, and the apparent tilt at each time window was expressed as the angle between this vector and that obtained for the pre-injection structure. Across the series, the His175 imidazole ring displayed an apparent progressive reorientation, reaching a displacement of approximately 0.8 Å and a tilt of approximately 20° relative to the pre-injection structure (Fig. S15). Because the same crystal volume was repeatedly irradiated during the experiment, we cannot exclude a contribution of radiation damage to this gradual coordinate shift observed only in this TR-FRX series. We therefore do not interpret this observation as direct evidence of a catalysis-associated motion but report it here as a structural change whose origin cannot currently be assigned unambiguously to either catalysis or radiation damage or both.

### *F*_o (t)_ − *F*_o (t0)_ isomorphous difference maps calculation

*F*_obs (t)_ − *F*_obs (t0)_ difference maps were computed using *phenix.fobs_minus_fobs_map* by pairing each dataset MTZ against a reference MTZ (subdataset before injection). For each pair, observed structure factors were taken from the same amplitude/sigma columns in both MTZ files, and model phases were calculated from a reference PDB. Maps were calculated with a 10 Å low-resolution cutoff. The high-resolution cutoff was adjusted to each subdataset for individual-map analyses, but fixed to a common value across the series for difference-density integration. Only reflections with F ≥ 3σ were retained, multiscale scaling and figure-of-merit weighting were applied.

### *F*_o_− *F*_o_ isomorphous difference maps density integration and analysis

For each region of interest, a mask was built as the union of spheres centered on every non-hydrogen atom (radii of 2 Å for loop conformations, 1.6 Å for OAA and 1.5 Å for GlcNAc), with shared volume counted once. Integration summed every voxel inside the mask (positive and negative) and multiplied by the voxel volume, with no σ cutoff applied. Timepoints are the midpoints of each acquisition window, since each integral is averaged over its time window (Fig. S13 A-C). For every map and every mask, the same mask shape was placed at 300 random positions in bulk solvent, rejecting any placement contacting the protein envelope (within 5 Å of modelled atoms). The standard deviation of these controls was taken as the error bar for the corresponding point (Fig. S13 D-F). Time constants are extracted from weighted (1/σ²) least-squares fits of y = A(1 − e^(−t/τ)), with no free offset (fixed origin). Confidence intervals were obtained by parametric bootstrap: each point was resampled within its error bar, and the model refitted. This procedure was repeated N = 2000 times and the interval limits are extracted from the 2.5th and 97.5th percentiles of the resulting distribution (Fig. S13 G-I). Integrated densities are reported in map·Å³, the sum of difference-map values within the mask multiplied by the voxel volume.

---

**Figure S1.**
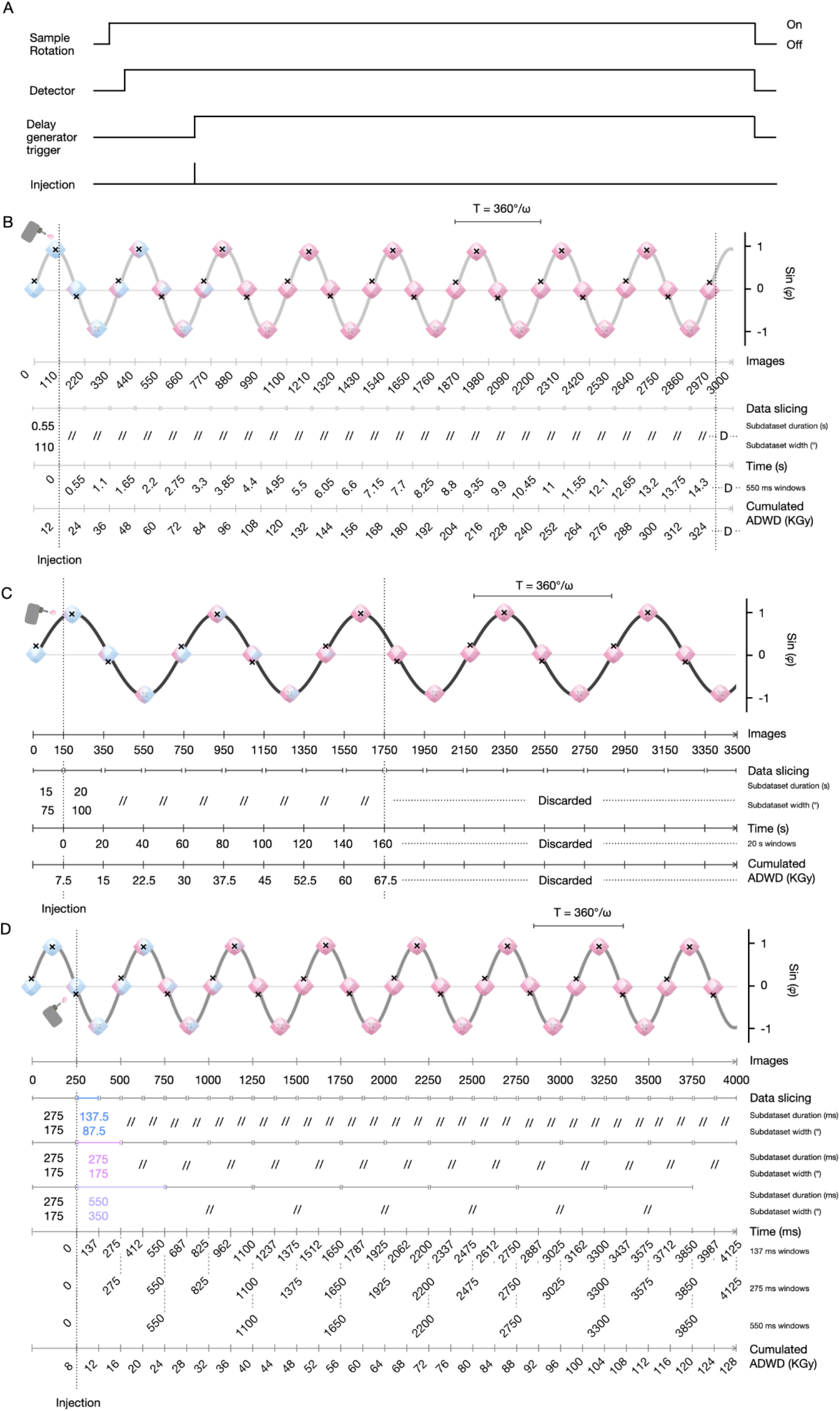
Schematic on TR-FRX triggering, data acquisition and subdataset partitioning. (A) Timing diagram describing the activation sequence of the main TR-FRX elements. Continuous sample rotation starts first, followed by detector acquisition. A duplicate signal from the detector is rooted through the delay generator to activate the nanodispenser after a controlled/applied delay. (B–D) Schematics of crystal rotation and partitioning for the three data collections. In each panel the trace (sin φ) is a schematic representation of the continuous rotation, where φ = ωt is the rotation angle of the crystal around the goniometer axis and ω the angular velocity. The crystal icons mark the orientation at successive points of the rotation, the black cross indicating a fixed point on the crystal. The color of the icons describes progressive ligand diffusion into the crystal, from blue (before injection) to pink (end of diffusion). The nanodispenser icon indicates the moment of ligand injection onto the crystal. T = 360°/ω represents the rotation period. Below each trace are indicated: image number; data partitioning (wedges duration and width, double slashes indicate repetition of the preceding value), the corresponding time windows and the cumulated average diffraction-weighted dose (ADWD). Subdatasets labelled Discarded or D did not yield complete, correctly indexed subdatasets and were excluded from further analysis. (B) HEWL acquisition: 3000 images at 1° and 5 ms per image (200 °/s), injection after image 110, sliced into 110-image subdatasets of 0.55 s and 110°. (C) CaMDH slow TR-FRX acquisition: 3500 images at 0.5° and 100 ms per image (5 °/s), injection after image 150, the reference subdataset spans 15 s and 75°, subsequent subdatasets 20 s and 100°. (D) CaMDH fast TR-FRX acquisition: 4000 images at 0.7° and 1.1 ms per image (636.4 °/s), injection after image 250; the same data were partitioned into 137, 275 and 550 ms subdatasets (87.5, 175 and 350°), shown in blue, purple and light purple.

**Figure S2.**
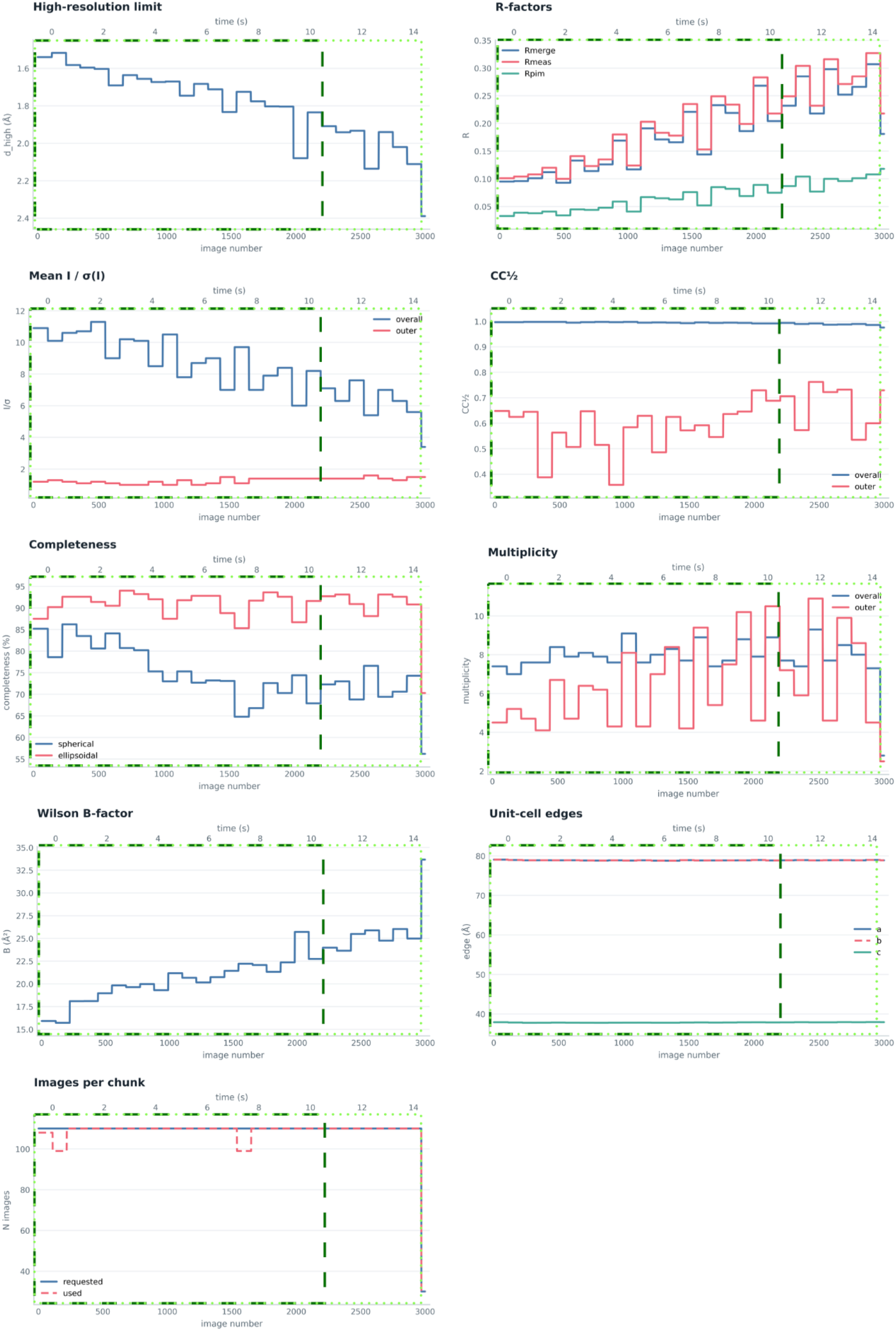
Evolution of key radiation-damage data-quality metrics during TR-FRX experiment on HEWL. Dark green dotted boxes indicate the subdatasets used for structure refinement. Light green boxes indicate the subdatasets used in initial Fo-Fo analysis.

**Figure S3.**
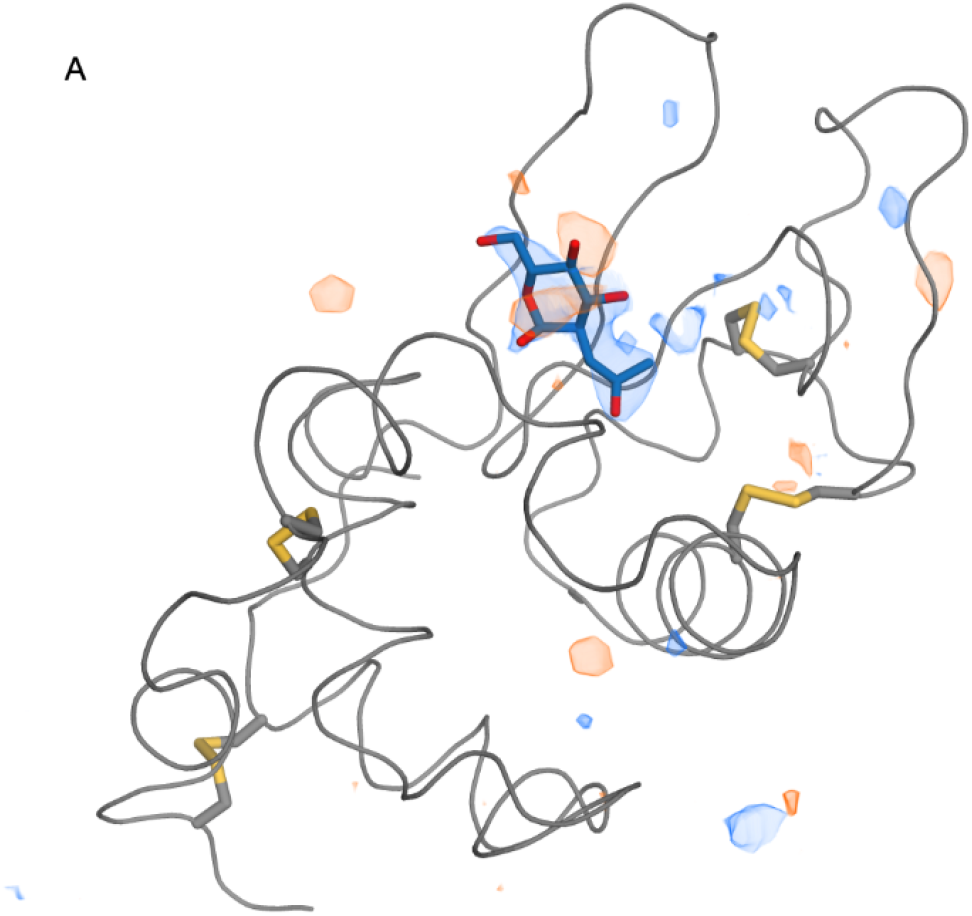
Fo_(11 s)_ − Fo_(apo)_ difference map across the HEWL asymmetric unit. Difference-density features are concentrated in the ligand binding cleft, strong positive peak (blue) corresponding to GlcNAc (dark blue, sticks) binding. Maps are contoured at ± 3.5 σ, with positive density in blue and negative density in orange.

**Figure S4.**
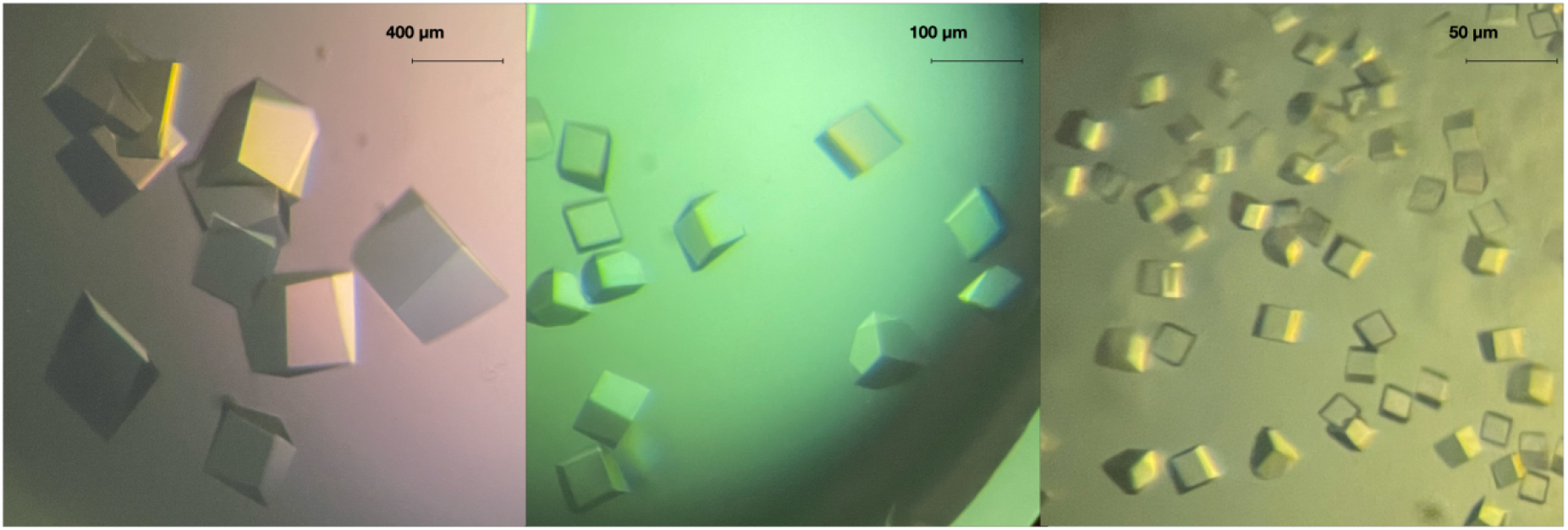
CaMDH crystals.

**Figure S5.**
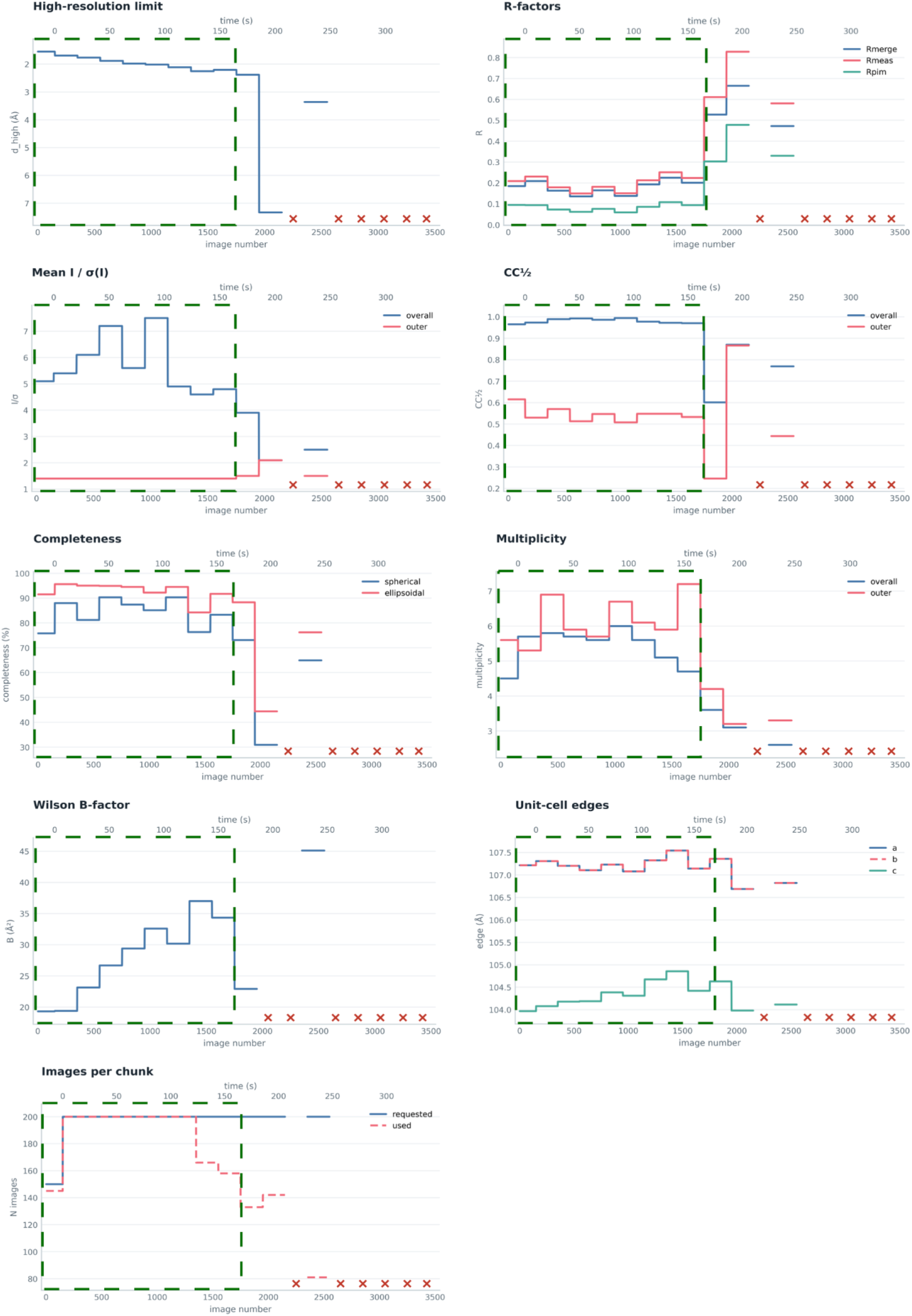
Evolution of key radiation-damage data-quality metrics during the slow-regime CaMDH TR-FRX experiment. Dark green boxes indicate the subdatasets used for structures refinement.

**Figure S6.**
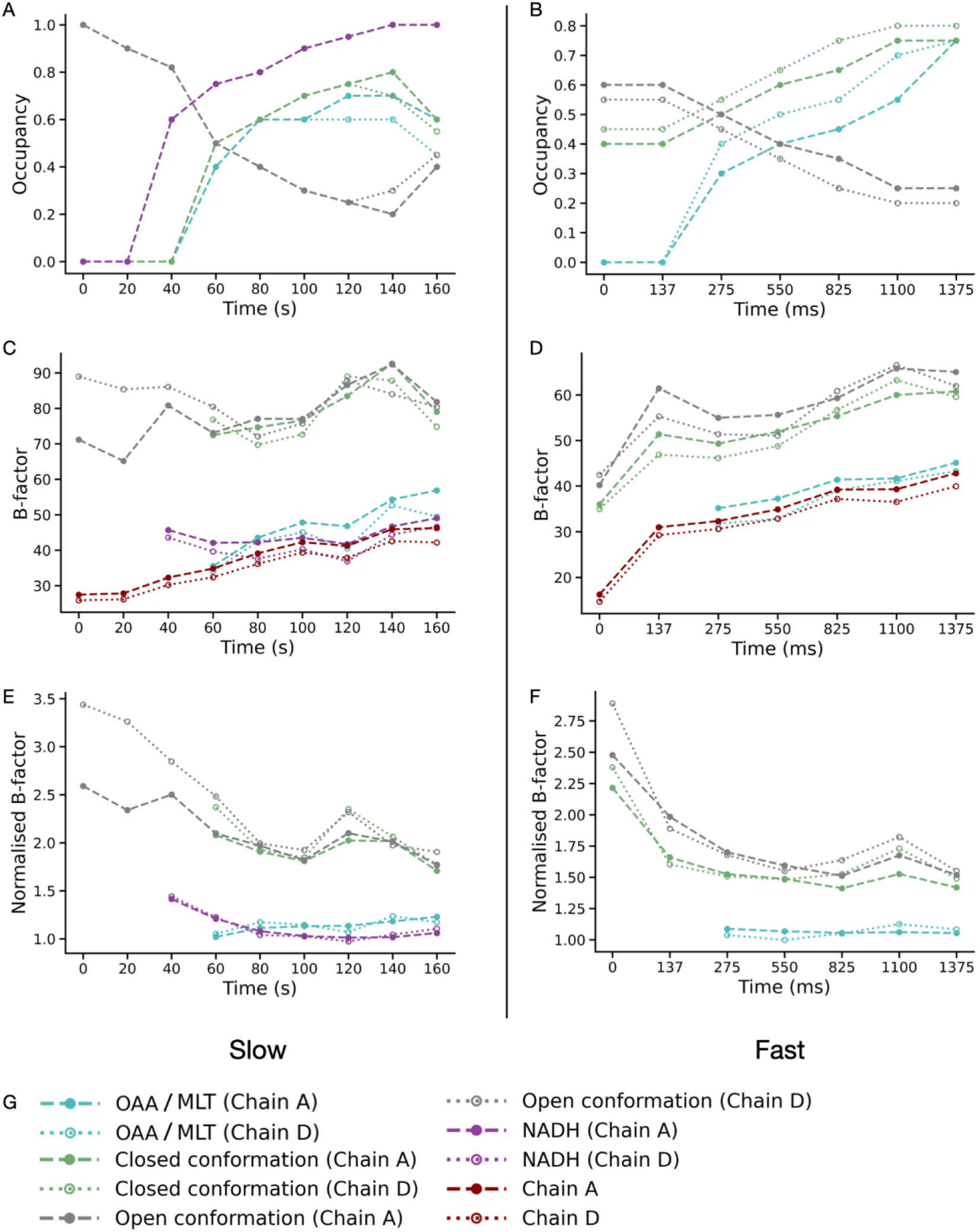
Occupancy and B-factor evolution of refined catalytic components for both slow and fast TR-FRX experiments. (A, **B)** Time-dependent evolution of refined occupancies for key catalytic components, including substrate/product (MLT and OAA), coenzyme (NADH), and active-site loop conformations open and closed **(C, D)** Time evolution of average B-factors (Å²) for key catalytic elements shown respectively for the slow and fast experiments **(E, F)** Time evolution of normalised B-factors (Å²) for key catalytic elements. **(G)** Legend of plotted elements.

**Figure S7.**
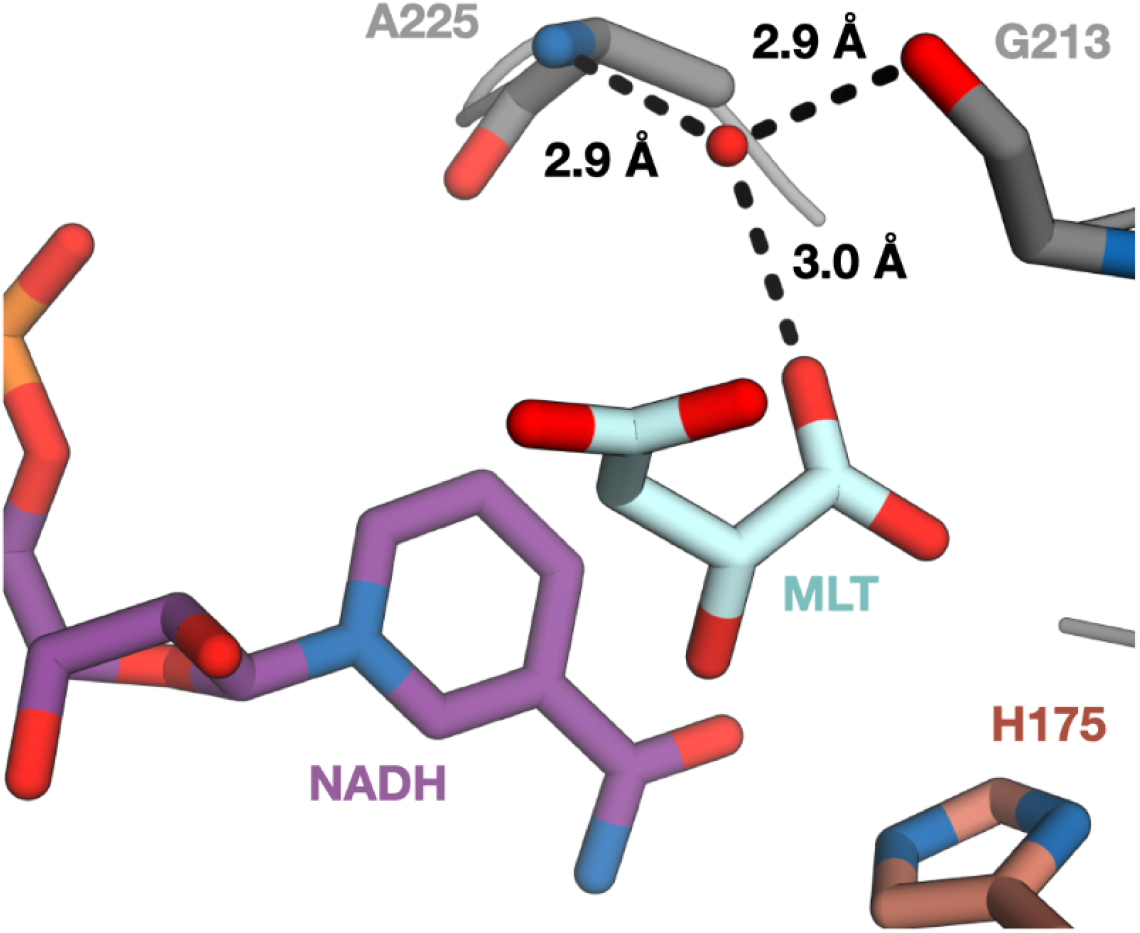
Hydrogen-bonding network stabilising a conserved water molecule in interaction with substrate. A water molecule (red sphere) mediates interactions with nearby residues A225 and G213 and the substrate (cyan). The key catalytic elements are displayed in stick representation. Dashed lines indicate hydrogen bonds.

**Figure S8.**
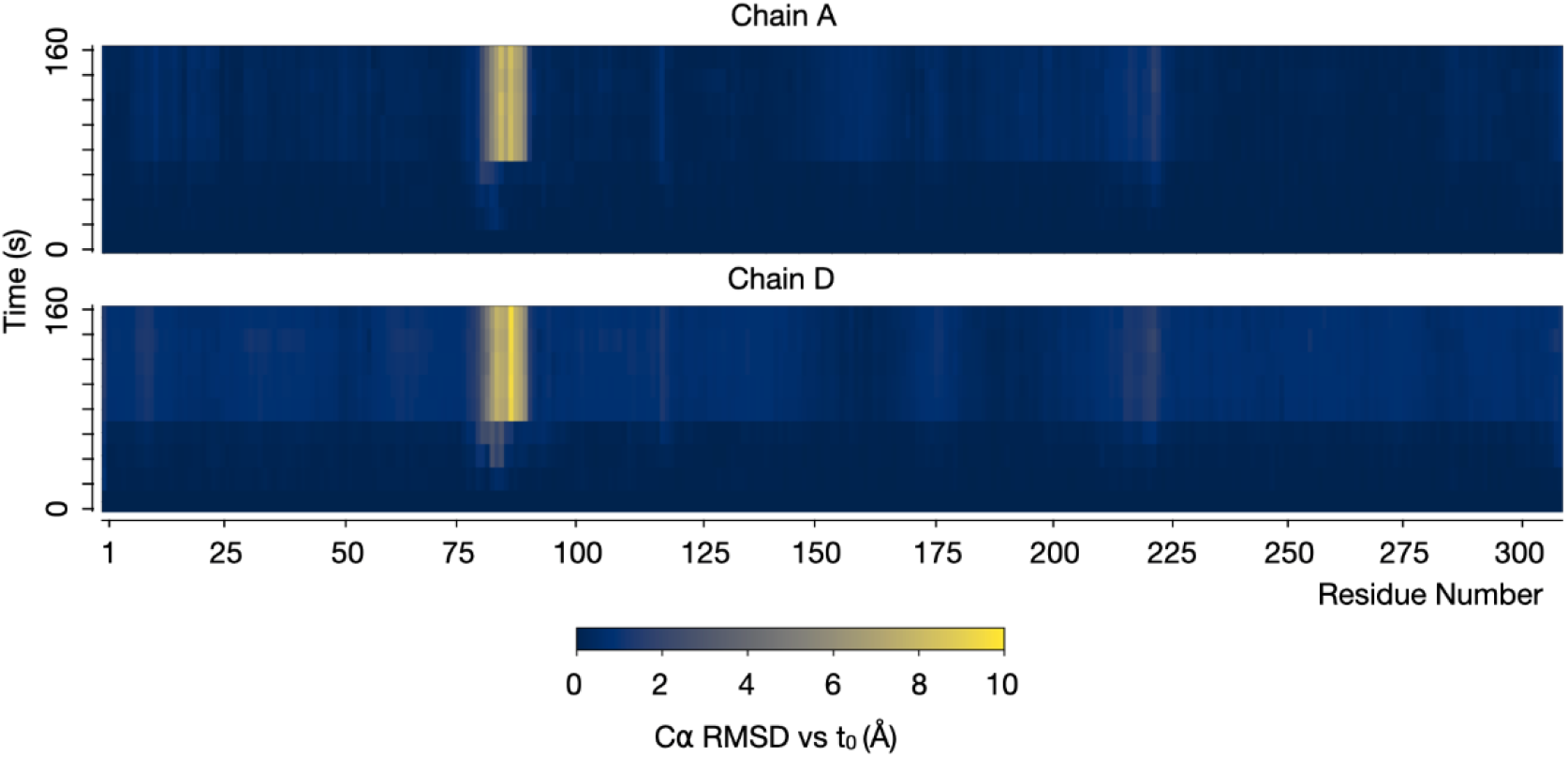
Time-dependent evolution of per-residue RMSD of chains A (top) and D (bottom) from CaMDH in the seconds to minutes TR-FRX series. Heatmaps showing the evolution of C⍺ RMSD (Å) relative to the reference structure at t=0. Color intensity encodes the magnitude of structural deviation, ranging from low (dark blue, 0 Å) to high (yellow, up to 10 Å).

**Figure S9.**
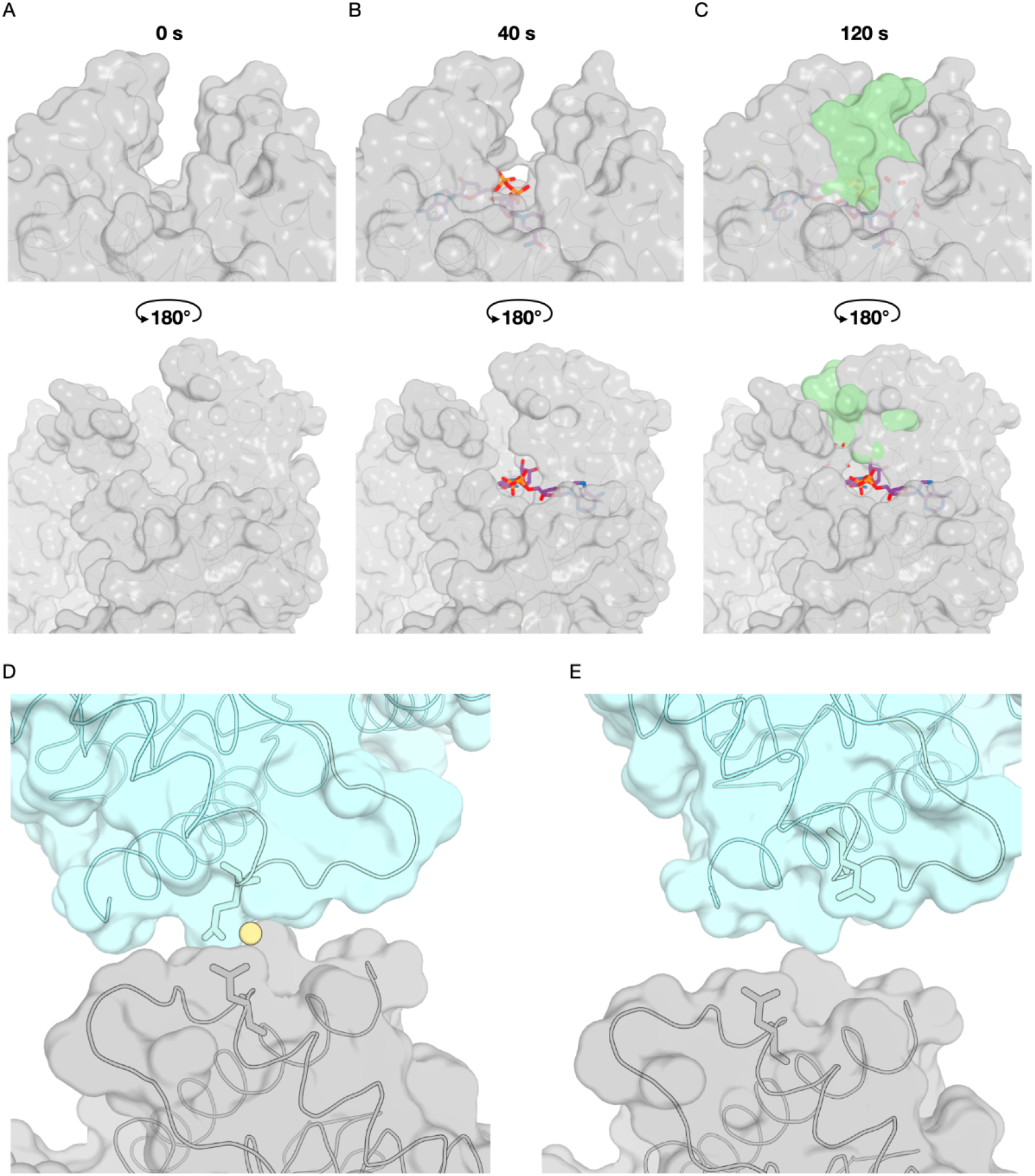
Catalytic loop dynamics and Cd²⁺-mediated crystal contact. (A–C) Views of the active site at (A) 0 s, (B) 20 - 40 s, and (C) 100 - 120 s after injection of a mixture containing 50 mM NADH and 50 mM OAA. MLT and NADH are shown as cyan and purple sticks, respectively. Closure of the catalytic loop (pale green surface) generates a fully sealed active-site pocket. (D) in the initial subdataset, a Cd²⁺ ion (light yellow sphere) bridges Glu89 and Asp90 from chain A to symmetry-related side chains (cyan surface). (E) Absence of the Cd²⁺-mediated crystal contact in chain D. The original molecule is shown as a grey surface, and the symmetry-related molecule as a cyan surface.

**Figure S10.**
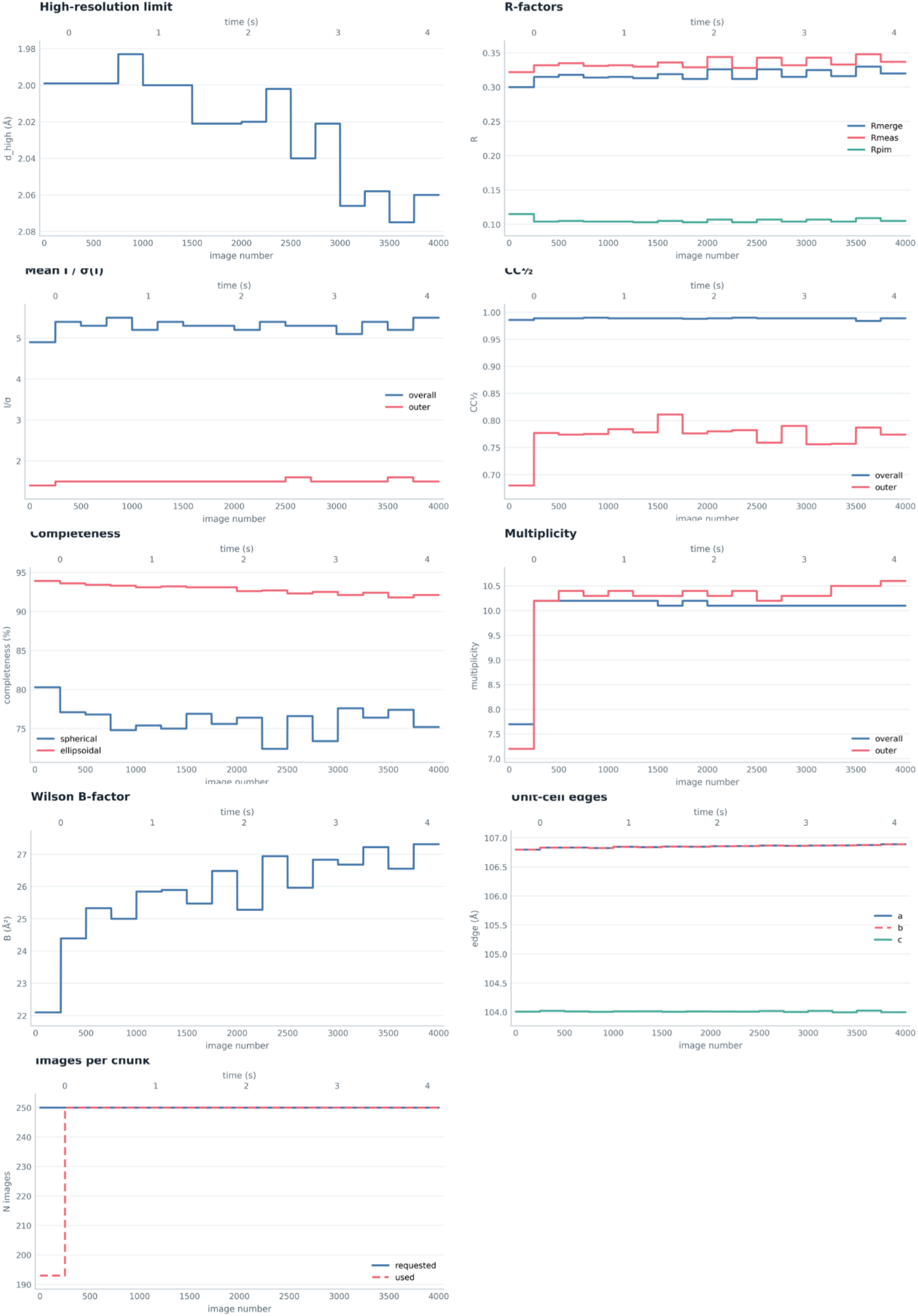
Evolution of key radiation-damage data-quality metrics during the fast-regime CaMDH TR-FRX control experiment.

**Figure S11.**
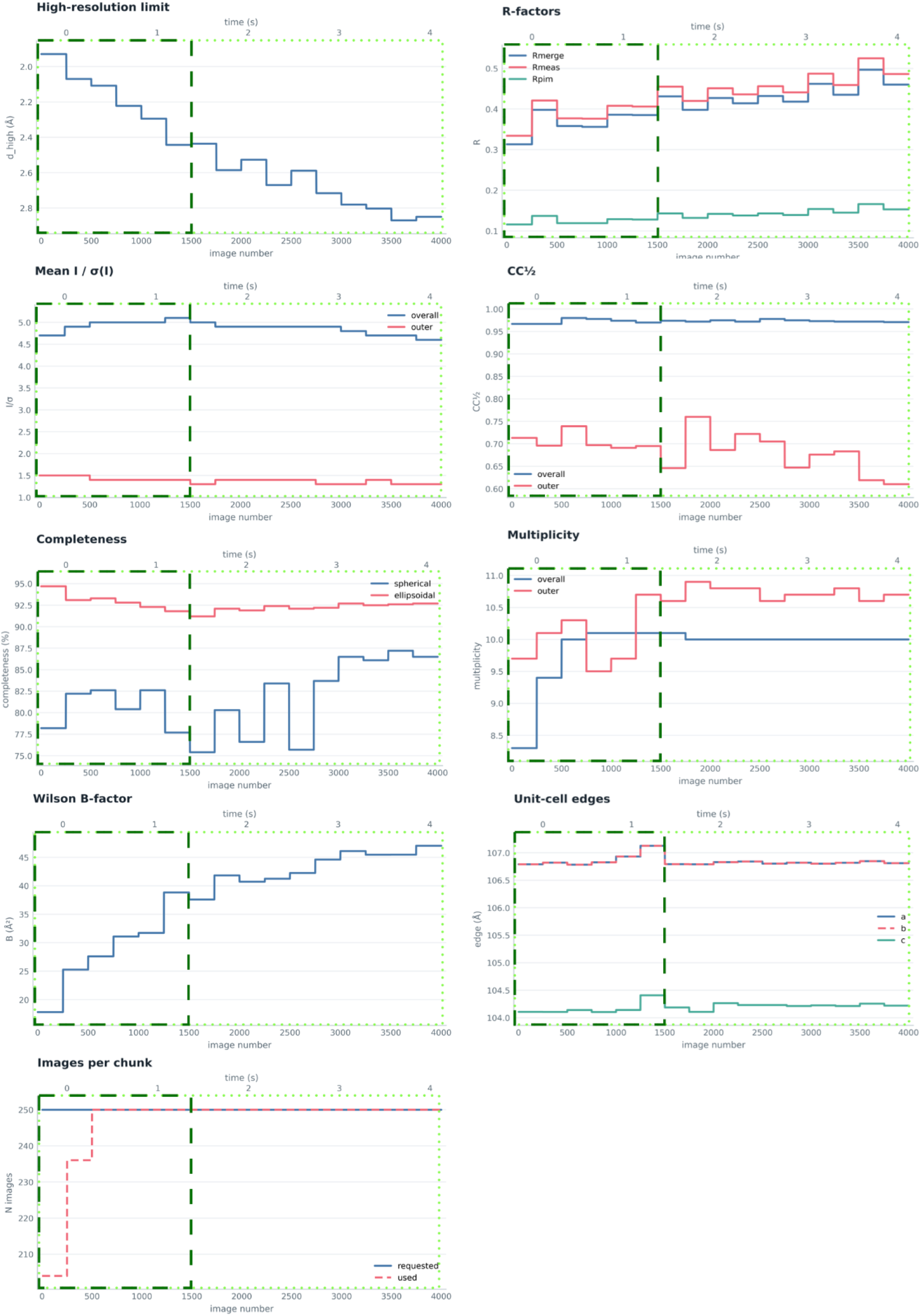
Evolution of key radiation-damage data-quality metrics during the fast-regime CaMDH TR-FRX experiment. Dark green boxes indicate the subdatasets used for structures refinement. Light green boxes indicate the subdatasets used for initial difference maps analysis.

**Figure S12.**
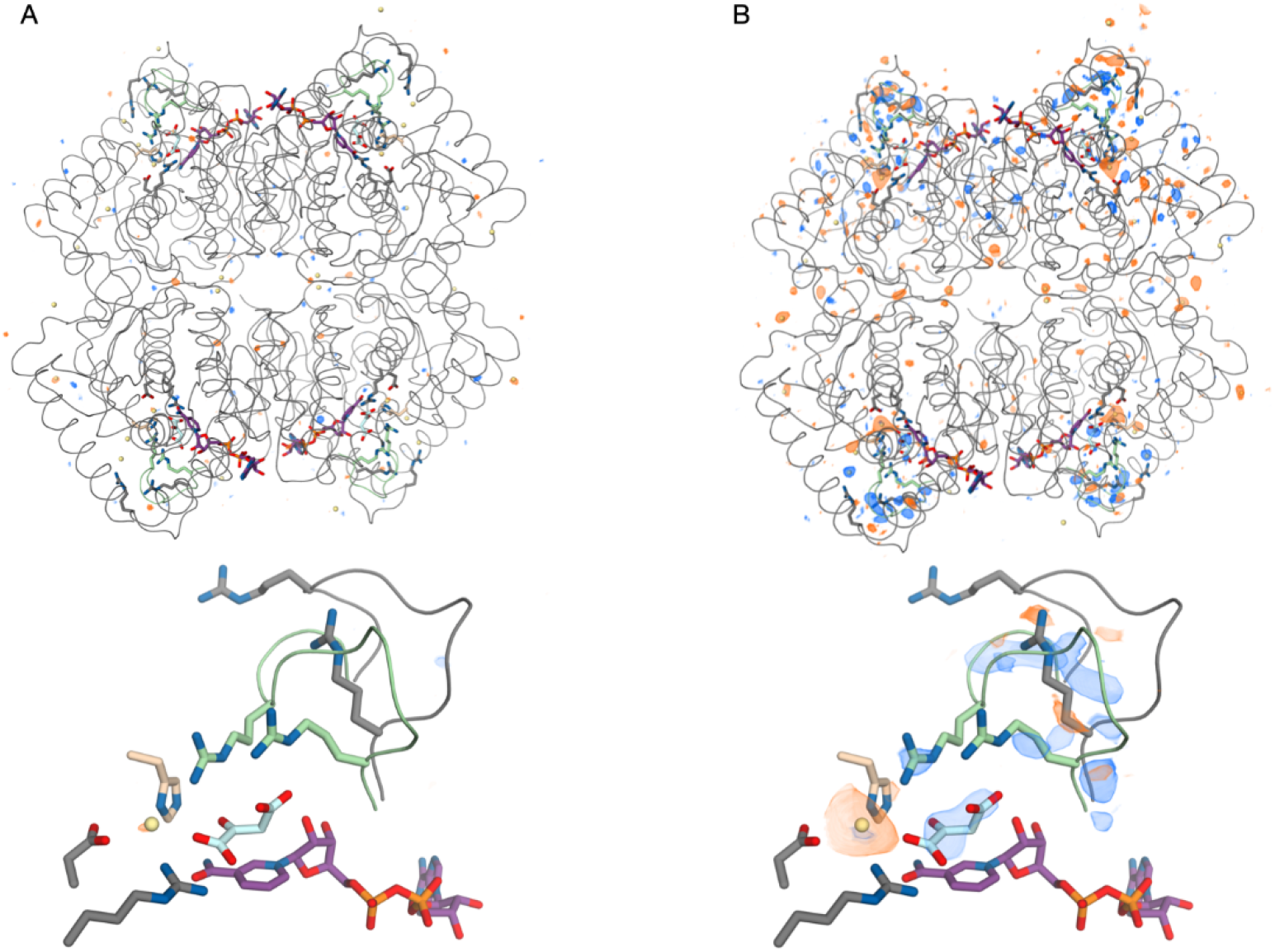
F_o_ − F_o_ difference maps of CaMDH with and without substrate injection. (A) F_o (1650 ms)_ − F_o(pre-injection)._ calculated for the control dataset without injection. (B) F_o (1650 ms)_ − F_o(pre-injection_ calculated for the dataset after injection. Positive density is shown in blue, negative in orange. All maps are contoured at ±4 σ. Top panels show the tetrameric assembly, bottom panels the active site (with NCS maps), with the mobile loop (residues 81–91) in grey (open loop) and green (closed loop), OAA in cyan, NAD⁺ in purple. A low occupancy Cd^2+^ atom present close to the active site is presented as a yellow sphere.

**Figure S13.**
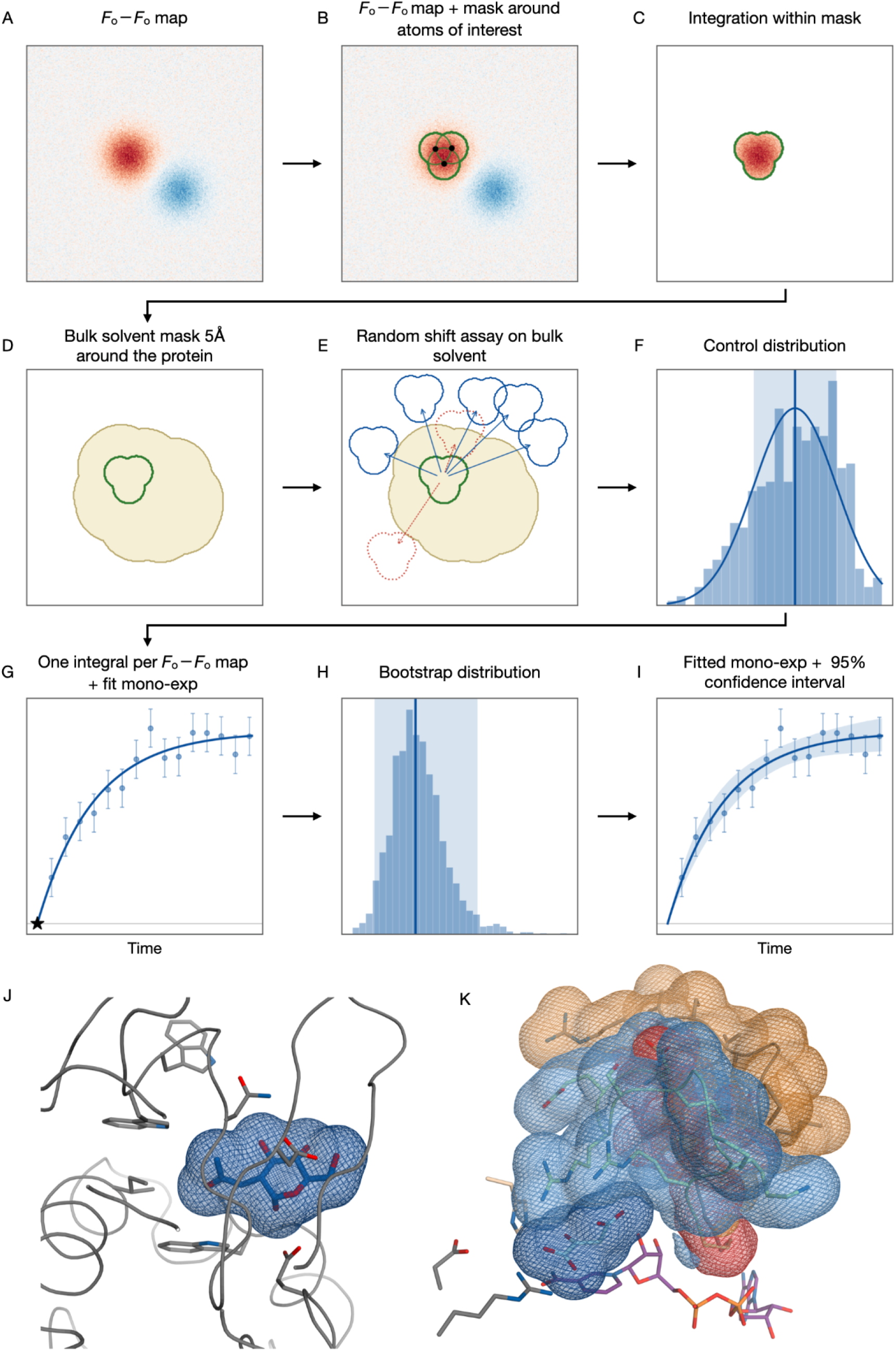
F_o_–F_o_ density integration. (A) F_o_**−**F_o_ density maps, appearing density in blue and leaving density in orange. (B) A mask is defined as a union of spheres (green outline) of radius r = 2 Å around every non hydrogen atoms of the selected residues, in which any shared volume is counted once. (C) All voxels within the mask are summed and multiplied by the voxel volume giving one number per map, with no σ threshold cutoff applied. (D) An exclusion envelope is defined 5 Å around every modelled atom (yellow); bulk solvent is everything outside it. (E) The integration mask is translated randomly to solvent positions 300 times (blue outline), any position overlapping the envelope (red outline) is rejected and redrawn. (F) The resulting distribution of N = 300 control integrals gives a mean, subtracted from the integrated density as a baseline, and a standard deviation, used as an error bar. (G) A time series is built from the integral of each unique map, and a weighted mono-exponential is fitted, forced through the origin (star), with each point placed at the midpoint of its acquisition time window. (H) Each point is resampled within its own error bar, and the mono-exponential refitted N = 2000 times, the band obtained represents the 2.5-97.5 percentiles of the resulting time constants. (I) The same fit is traced with the 95% confidence interval band (blue shade) (J) Integration mask applied 1.5 Å around the GlcNAc binding site in HEWL. (K) Integration masks applied 2 Å around both conformations of the mobile loop, open (orange), closed (blue), overlapping and not integrated voxels shared by both alternate conformations (red), and mask 1.6 Å around ligand (dark blue) in CaMDH.

**Figure S14.**
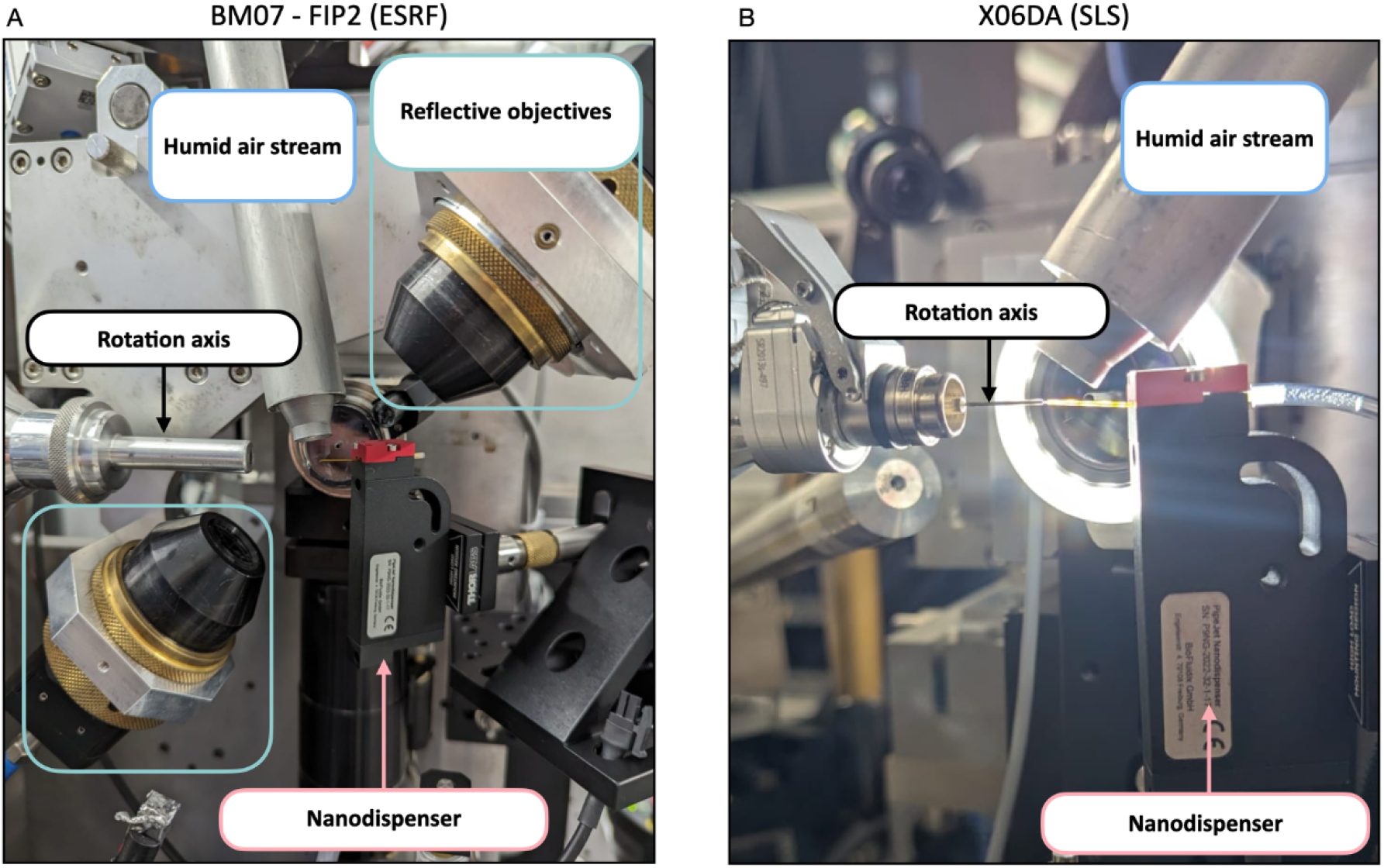
TR-FRX experimental set-up installed on the BM07 beamline at the ESRF (A) and on the X06DA beamline at the SLS 2.0 (B). The nanodispenser was positioned within the sample environment using a Thorlabs magnetic base mounted on manual translation stages, allowing adjustment along the X, Y, and Z axes relative to the mounted crystal. Initial tests were performed on the beamline BM07. The portability and rapid installation of the setup were subsequently demonstrated on the beamline X06DA during a single three-shift beamtime in November 2025. During this beamtime, the nanodispenser was installed for the first time in the sample environment.

**Figure S15.**
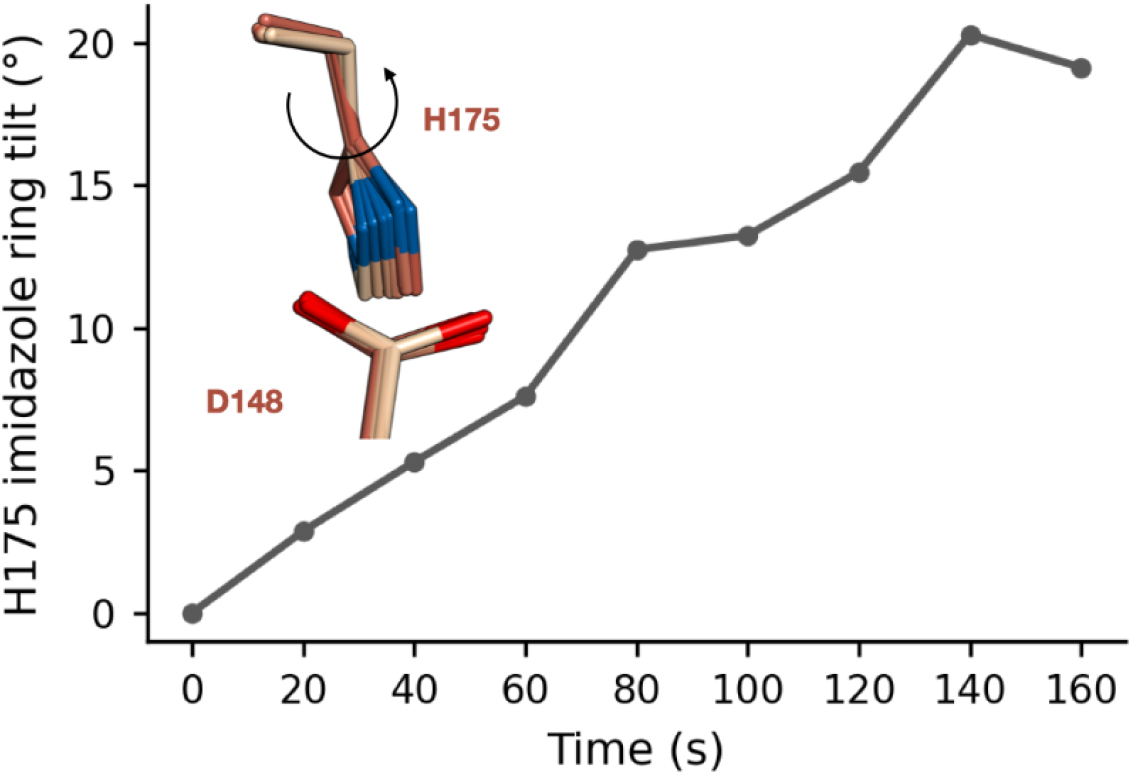
Progressive tilting of Histidine 175 during the course of the acquisition.

**Table S1.**
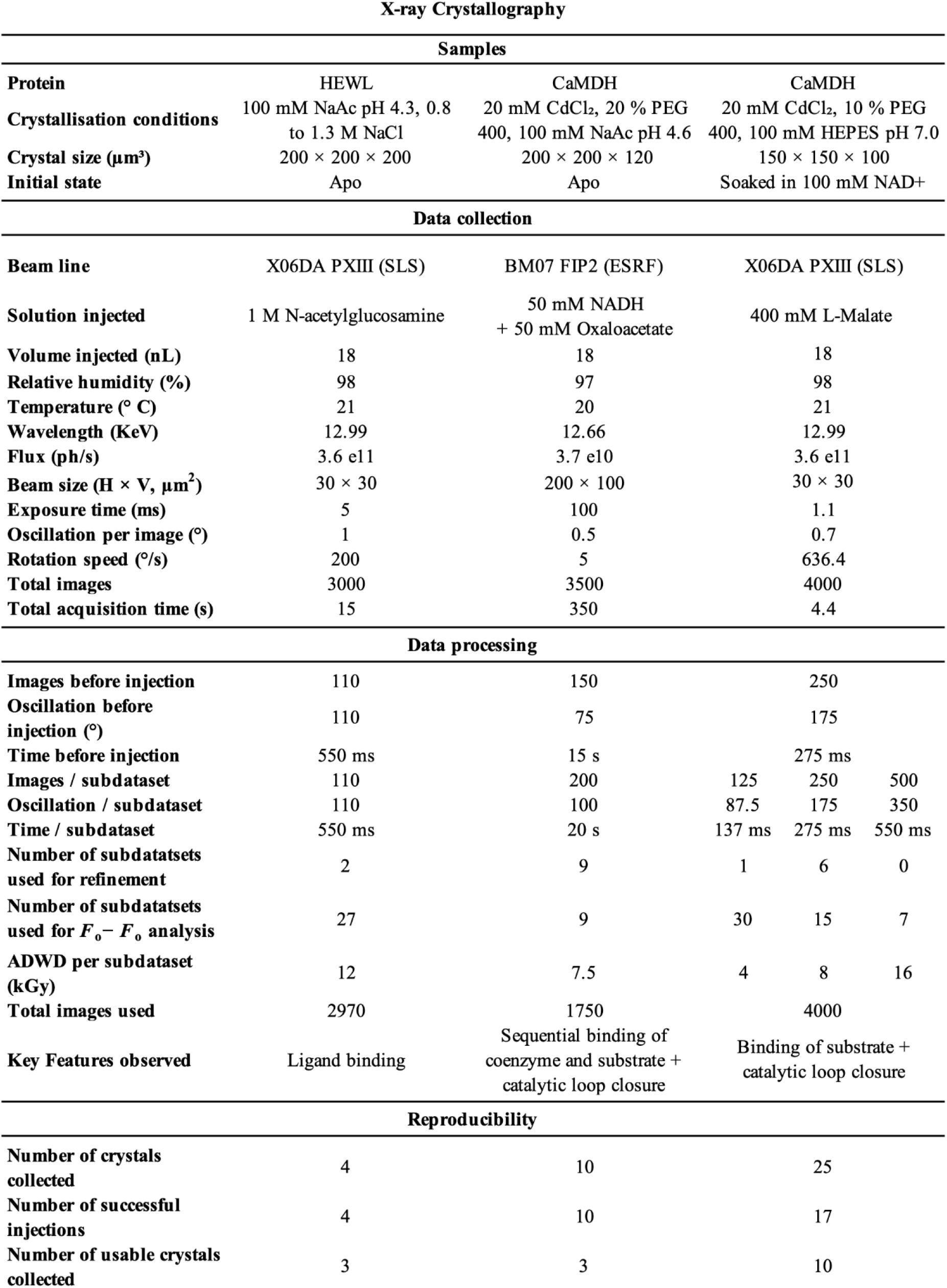
Summary of TR-FRX experiments performed on HEWL and CaMDH. For each experiment, the table reports the sample and reaction conditions, data-collection parameters, data-processing strategy, and reproducibility of the resulting structural series.

**Table S2.** Crystallographic data collection statistics of HEWL structures obtained before ligand injection (apo) and at 10.45 seconds following the injection of 1M GlcNAc. Numbers in parentheses denote the highest resolution shell.

|  | HEWL Apo | HEWL 10.45s |
| --- | --- | --- |
| Space group | $P4_32_12$ | |
|  | 79.067 | 78.848 |
| $a, b, c$ (Å) | 79.067 | 78.848 |
|  | 37.905 | 37.864 |
| Resolution range (Å) | 55.91 - 1.54 | 55.75 - 1.83 |
|  | (1.63 - 1.54) | (1.95 - 1.83) |
| $R_{\text{pim}}$ | 0.033 (0.480) | 0.075 (0.669) |
| $CC_{1/2}$ | 0.997 (0.647) | 0.992 (0.689) |
| $I/\sigma(I)$ | 10.9 (1.2) | 8.2 (1.4) |
| Ellipsoidal |  |  |
| Completeness (%) | 87.9 (32.7) | 91.6 (57.7) |
| Multiplicity | 7.4 (4.5) | 8.9 (10.5) |
| Total reflections | 116233 (3493) | 66141 (3900) |
| Unique reflections | 15622 (781) | 7422 (371) |
| $R_{\text{work}}/R_{\text{free}}$ | 0.1575 / 0.1785 | 0.1615 / 0.1951 |
| No. atoms | 1169 | 1163 |
| Macromolecules | 1070 | 1070 |
| Ligands | 3 | 18 |
| Solvent | 96 | 75 |
| B-factors (Å<sup>2</sup>) | 27.22 | 36.50 |
| Macromolecules | 26.22 | 36.15 |
| Ligands | 37.93 | 37.77 |
| Solvent | 37.98 | 41.22 |
| R.m.s. deviations |  |  |
| Bond lengths (Å) | 0.008 | 0.006 |
| Bond angles (°) | 0.93 | 0.85 |
| PDB code | 29PU | 29PV |

**Table S3.** Diffusion times (ms) required to reach 10, 25, 50, 75, and 99% of the external ligand concentration at the center of parallelepipedic crystals of varying dimensions. Values are computed using Eq. 2 from Schmidt, Crystals 2020, using the diffusion coeflcient D = 2.3 × 10⁻⁶ cm²/s. Reducing edge length from 200 µm to 30 µm decreases diffusion times by more than two orders of magnitude.

| Saturation (%) | 200×200×200 $\mu\text{m}$ | 200×200×100 $\mu\text{m}$ | 30×30×100 $\mu\text{m}$ | 30×30×30 $\mu\text{m}$ |
| --- | --- | --- | --- | --- |
| 10% | 3834 | 1413 | 98 | 86 |
| 25% | 5446 | 2273 | 146 | 123 |
| 50% | 8184 | 3812 | 232 | 184 |
| 75% | 12378 | 6074 | 370 | 279 |
| 99% | 31306 | 15651 | 1001 | 704 |

**Table S4.**
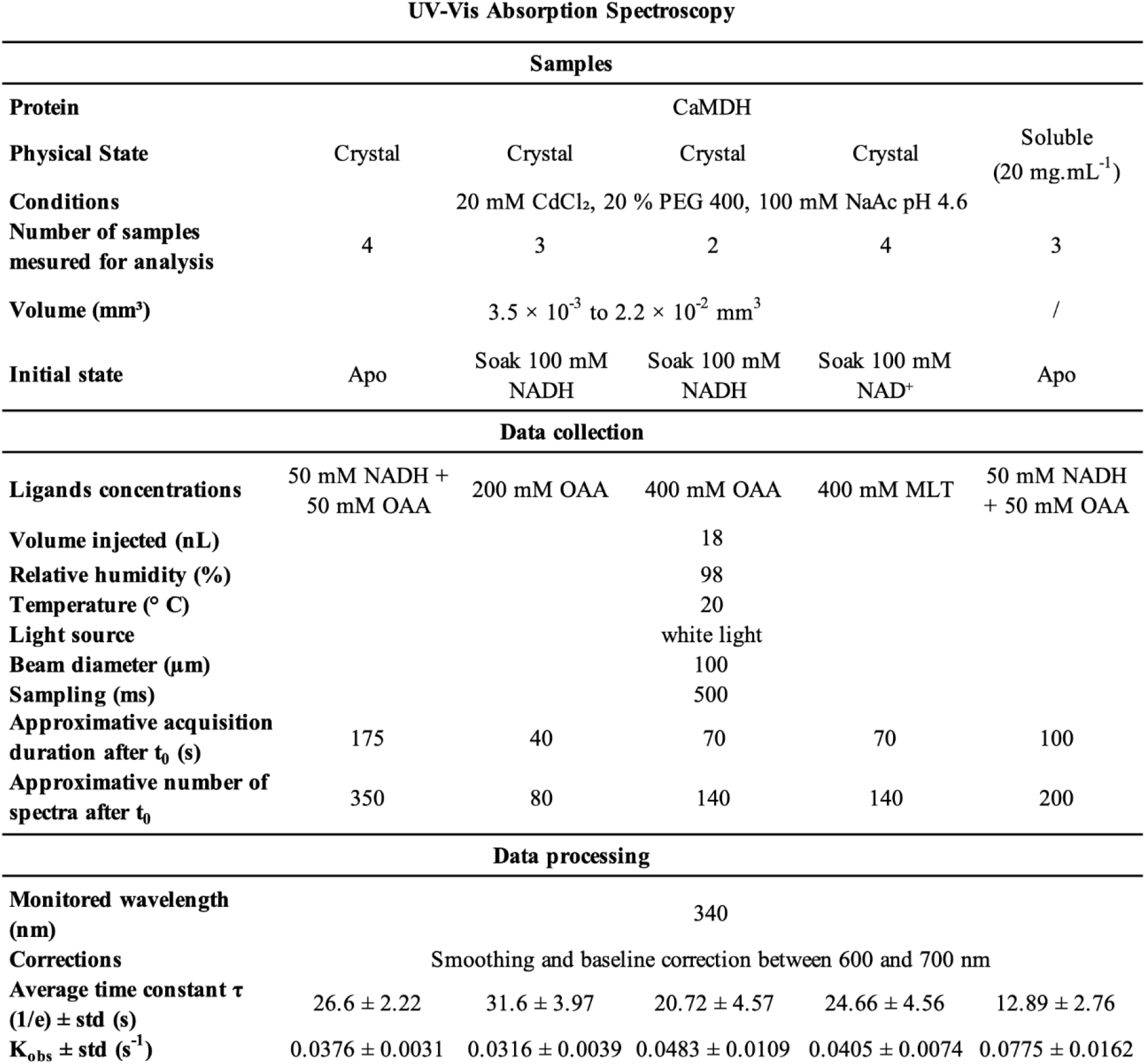
Summary of UV-Vis absorption spectroscopy measurements on CaMDH. For each experiment, the table reports the sample conditions, ligand-delivery parameters, data acquisition settings, and data processing results. Apparent time constants (tau) were obtained from mono-exponential fits to the kinetic traces monitored at 340 nm and are reported as averages across independent samples. T_0_ corresponds to the injection which triggers the in crystallo reaction. Apparent observed rate constants (K_obs_) were calculated as 1/tau and are provided for descriptive comparison between experimental conditions (see method section in supporting information for details).

**Table S5.** Crystallographic data collection and refinement statistics for the series of CaMDH structures obtained prior to ligand injection (apo) and at 20-second intervals following the injection of an equimolar mixture of NADH and OAA (50 mM). Numbers in parentheses denote the highest resolution shell.

|  | *CaMDH apo* | *CaMDH 20 s* | *CaMDH 40 s* | *CaMDH 60 s* | *CaMDH 80 s* |
| --- | --- | --- | --- | --- | --- |
| Space group | *P*3<sub>1</sub>21 |  |  |  |  |
| *a*, *b*, *c* (Å) | 107.217 | 107.307 | 107.202 | 107.107 | 107.235 |
|  | 107.217 | 107.307 | 107.202 | 107.107 | 107.235 |
|  | 103.968 | 104.079 | 104.179 | 104.189 | 104.390 |
| Resolution range (Å) | 92.85 - 1.55 | 92.93 - 1.69 | 92.84 - 1.76 | 92.76 - 1.88 | 92.87 - 1.98 |
|  | (1.66 - 1.55) | (1.79 - 1.69) | (1.89 - 1.76) | (1.97 - 1.88) | (2.10 - 1.98) |
| R<sub>pim</sub> | 0.095 (0.525) | 0.094 (0.602) | 0.073 (0.556) | 0.062 (0.559) | 0.076 (0.543) |
| CC<sub>1/2</sub> | 0.965 (0.615) | 0.973 (0.530) | 0.989 (0.570) | 0.992 (0.515) | 0.987 (0.547) |
| I/σ(I) | 5.1 (1.4) | 5.4 (1.4) | 6.1 (1.4) | 7.2 (1.4) | 5.6 (1.4) |
| Ellipsoidal Completeness (%) | 91.5 (62.9) | 95.6 (52.0) | 94.9 (59.7) | 95.0 (51.0) | 94.6 (47.2) |
| Multiplicity | 4.5 (5.6) | 5.7 (5.3) | 5.8 (6.9) | 5.7 (5.9) | 5.6 (5.7) |
| Total reflections | 342283 (21210) | 389916 (18199) | 323080 (19129) | 290726 (14932) | 238569 (12035) |
| Unique reflections | 75923 (3796) | 68295 (3115) | 55553 (2778) | 50972 (2549) | 42600 (2130) |
| R<sub>work</sub>/R<sub>free</sub> | 0.1593 / 0.1804 | 0.1503 / 0.1893 | 0.1570 / 0.1892 | 0.1476 / 0.1827 | 0.1572 / 0.1901 |
| No. atoms | 5148 | 5152 | 5073 | 5236 | 5207 |
| Macromolecules | 4713 | 4702 | 4650 | 4833 | 4828 |
| Ligands | 9 | 9 | 97 | 115 | 115 |
| Solvent | 426 | 441 | 326 | 288 | 264 |
| B-factors (Å<sup>2</sup>) | 29.57 | 29.98 | 33.77 | 37.00 | 40.59 |
| Macromolecules | 28.66 | 28.81 | 32.98 | 36.53 | 40.17 |
| Ligands | 39.99 | 44.96 | 44.94 | 40.76 | 41.77 |
| Solvent | 39.34 | 42.17 | 41.77 | 43.27 | 47.74 |
| R.m.s. deviations |  |  |  |  |  |
| Bond lengths (Å) | 0.011 | 0.014 | 0.007 | 0.009 | 0.004 |
| Bond angles (°) | 1.05 | 1.23 | 0.86 | 0.98 | 0.68 |
| PDB code | 29PE | 29PF | 29PG | 29PH | 29PI |

**Table S5.** Crystallographic data collection and refinement statistics for the series of *CaMDH* structures obtained prior to ligand injection (*apo*) and at 20-second intervals following the injection of an equimolar mixture of NADH and OAA (50 mM).
|  | *CaMDH 100 s* | *CaMDH 120 s* | *CaMDH 140 s* | *CaMDH 160 s* |
| --- | --- | --- | --- | --- |
| Space group | *P*3<sub>1</sub>21 |  |  |  |
| *a*, *b*, *c* (Å) | 107.082 | 107.325 | 107.535 | 107.147 |
|  | 107.082 | 107.325 | 107.535 | 107.147 |
|  | 104.313 | 104.673 | 104.848 | 104.423 |
| Resolution range (Å) | 92.74 - 2.02 | 69.50 - 2.11 | 93.13 - 2.25 | 69.36 - 2.20 |
|  | (2.13 - 2.02) | (2.21 - 2.11) | (2.38 - 2.25) | (2.34 - 2.20) |
| R<sub>pim</sub> | 0.059 (0.624) | 0.086 (0.565) | 0.108 (0.650) | 0.094 (0.567) |
| CC<sub>1/2</sub> | 0.994 (0.509) | 0.978 (0.554) | 0.972 (0.545) | 0.970 (0.536) |
| I/σ(I) | 7.5 (1.4) | 4.9 (1.4) | 4.6 (1.4) | 4.7 (1.4) |
| Ellipsoidal Completeness (%) | 92.2 (51.7) | 94.5 (49.3) | 84.2 (50.4) | 91.7 (57.4) |
| Multiplicity | 6.0 (6.7) | 5.6 (6.0) | 5.1 (5.9) | 4.7 (1.4) |
| Total reflections | 232926 (13080) | 204552 (11044) | 130833 (7596) | 137502 (10625) |
| Unique reflections | 39053 (1853) | 36544 (1827) | 25589 (1279) | 29422 (1471) |
| R<sub>work</sub>/R<sub>free</sub> | 0.1563 / 0.1862 | 0.1509 / 0.1897 | 0.1665 / 0.1984 | 0.1469 / 0.1890 |
| No. atoms | 5178 | 5157 | 5095 | 5128 |
| Macromolecules | 4817 | 4815 | 4820 | 4810 |
| Ligands | 114 | 117 | 115 | 114 |
| Solvent | 247 | 225 | 160 | 204 |
| B-factors (Å<sup>2</sup>) | 43.61 | 43.08 | 47.44 | 46.81 |
| Macromolecules | 43.31 | 42.99 | 47.47 | 46.63 |
| Ligands | 44.10 | 42.68 | 48.71 | 49.46 |
| Solvent | 49.24 | 45.33 | 45.55 | 49.58 |
| R.m.s. deviations |  |  |  |  |
| Bond lengths (Å) | 0.003 | 0.013 | 0.002 | 0.003 |
| Bond angles (°) | 0.6 | 1.03 | 0.53 | 0.58 |
| PDB code | 29PJ | 29PK | 29PL | 29PM |

**Table S6.** Crystallographic data collection statistics for the series of CaMDH structures obtained prior to ligand injection (apo) and at 137 ms intervals following the injection of 400 mM MLT on presoaked crystal with 100 mM NAD^+^. Numbers in parentheses denote the highest resolution shell.

|  | *CaMDH* apo | *CaMDH* 137 ms | *CaMDH* 275 ms | *CaMDH* 550 ms |
| --- | --- | --- | --- | --- |
| Space group | *P*3<sub>1</sub>21 |  |  |  |
| *a*, *b*, *c* (Å) | 106.791 | 106.842 | 106.827 | 106.782 |
|  | 106.791 | 106.842 | 106.827 | 106.782 |
|  | 104.107 | 104.136 | 104.112 | 104.137 |
| Resolution range (Å) | 92.48 - 1.92 | 92.53 - 2.15 | 92.51 - 2.07 | 92.48 - 2.11 |
|  | (2.06 - 1.92) | (2.37 - 2.15) | (2.22 - 2.07) | (2.25 - 2.11) |
| R<sub>pim</sub> | 0.145 (0.573) | 0.205 (0.594) | 0.137 (0.593) | 0.119 (0.622) |
| CC<sub>1/2</sub> | 0.944 (0.688) | 0.894 (0.570) | 0.966 (0.704) | 0.978 (0.736) |
| I/σ(I) | 4.8 (1.5) | 3.8 (1.4) | 4.9 (1.5) | 5.0 (1.4) |
| Ellipsoidal Completeness (%) | 94.8 (59.4) | 92.6 (41.5) | 93.2 (46.6) | 93.3 (47.5) |
| Multiplicity | 8.3 (9.7) | 4.4 (4.1) | 9.3 (10.0) | 10.0 (10.4) |
| Total reflections | 342099 (20115) | 127622 (5918) | 323902 (17329) | 330125 (17100) |
| Unique reflections | 41466 (2073) | 28795 (1440) | 34731 (1737) | 33047 (1652) |
| R<sub>work</sub> / R<sub>free</sub> | 0.1597 / 0.2068 | 0.1740 / 0.2256 | 0.1630 / 0.1993 | 0.1604 / 0.2004 |
| No. atoms | 5431 | 5248 | 5260 | 5297 |
| Macromolecules | 4869 | 4831 | 4848 | 4853 |
| Ligands | 108 | 110 | 128 | 128 |
| Solvent | 454 | 307 | 284 | 316 |
| B-factors (Å<sup>2</sup>) | 18.17 | 32.24 | 33.14 | 35.62 |
| Macromolecules | 17.17 | 31.87 | 32.87 | 35.21 |
| Ligands | 16.05 | 32.78 | 33.37 | 35.77 |
| Solvent | 29.35 | 37.83 | 37.63 | 41.93 |
| R.m.s. deviations |  |  |  |  |
| Bond lengths (Å) | 0.008 | 0.003 | 0.004 | 0.005 |
| Bond angles (°) | 0.88 | 0.60 | 0.70 | 0.68 |
| PDB code | 29PN | 29PO | 29PP | 29PQ |
|  | *CaMDH* 825 ms | *CaMDH* 1100 ms | *CaMDH* 1375 ms |  |
| Space group | *P*3<sub>1</sub>21 |  |  |  |
| *a*, *b*, *c* (Å) | 106.826 | 106.931 | 107.129 |  |
|  | 106.826 | 106.931 | 107.129 |  |
|  | 104.105 | 104.141 | 104.409 |  |
| Resolution range (Å) | 92.51 - 2.22 | 92.60 - 2.29 | 92.78 - 2.44 |  |
|  | (2.40 - 2.22) | (2.45 - 2.29) | (2.68 - 2.44) |  |
| R<sub>pim</sub> | 0.118 (0.629) | 0.128 (0.694) | 0.128 (0.640) |  |
| CC<sub>1/2</sub> | 0.979 (0.692) | 0.967 (0.669) | 0.969 (0.689) |  |
| I/σ(I) | 5.0 (1.4) | 5.0 (1.4) | 5.1 (1.4) |  |
| Ellipsoidal Completeness (%) | 92.8 (46.2) | 92.4 (44.7) | 91.9 (39.5) |  |
| Multiplicity | 10.1 (9.5) | 10.1 (9.7) | 10.1 (10.7) |  |
| Total reflections | 277112 (13081) | 259336 (12528) | 204288 (10852) |  |
| Unique reflections | 27544 (1377) | 25775 (1289) | 20279 (1014) |  |
| R<sub>work</sub> / R<sub>free</sub> | 0.1607 / 0.2007 | 0.1572 / 0.2167 | 0.1566 / 0.2106 |  |
| No. atoms | 5204 | 5244 | 5247 |  |
| Macromolecules | 4841 | 4867 | 4867 |  |
| Ligands | 128 | 128 | 128 |  |
| Solvent | 235 | 249 | 252 |  |
| B-factors (Å<sup>2</sup>) | 39.78 | 40.02 | 42.98 |  |
| Macromolecules | 39.67 | 39.84 | 42.89 |  |
| Ligands | 39.59 | 39.31 | 43.12 |  |
| Solvent | 42.19 | 44.01 | 44.80 |  |
| R.m.s. deviations |  |  |  |  |
| Bond lengths (Å) | 0.003 | 0.003 | 0.002 |  |
| Bond angles (°) | 0.56 | 0.55 | 0.52 |  |
| PDB code | 29PR | 29PS | 29PT |  |

## References

1. Henzler-Wildman KA, Thai V, Lei M, Ott M, Wolf-Watz M, Fenn T, et al. Intrinsic motions along an enzymatic reaction trajectory. Nature. 2007 Dec;450(7171):838–44. doi:10.1038/nature06410

2. Bar-Even A, Noor E, Savir Y, Liebermeister W, Davidi D, Tawfik DS, et al. The Moderately Efficient Enzyme: Evolutionary and Physicochemical Trends Shaping Enzyme Parameters. Biochemistry. 2011 May 31;50(21):4402–10. doi:10.1021/bi2002289

3. Brändén G, Neutze R. Advances and challenges in time-resolved macromolecular crystallography. Science. 2021 Aug 27;373(6558). doi:10.1126/science.aba0954

4. Fraser JS, Van Den Bedem H, Samelson AJ, Lang PT, Holton JM, Echols N, et al. Accessing protein conformational ensembles using room-temperature X-ray crystallography. Proc Natl Acad Sci. 2011 Sep 27;108(39):16247–52. doi:10.1073/pnas.1111325108

5. Jumper J, Evans R, Pritzel A, Green T, Figurnov M, Ronneberger O, et al. Highly accurate protein structure prediction with AlphaFold. Nature. 2021 Aug 26;596(7873):583–9. doi:10.1038/s41586-021-03819-2

6. Bourgeois D, Royant A. Advances in kinetic protein crystallography. Curr Opin Struct Biol. 2005 Oct;15(5):538–47. doi:10.1016/j.sbi.2005.08.002

7. Caramello N, Royant A. From femtoseconds to minutes: time-resolved macromolecular crystallography at XFELs and synchrotrons. Acta Crystallogr Sect Struct Biol. 2024 Feb 1;80(2):60–79. doi:10.1107/s2059798323011002

8. Aumonier S, Santoni G, Gotthard G, von Stetten D, Leonard GA, Royant A. Millisecond time-resolved serial oscillation crystallography of a blue-light photoreceptor at a synchrotron. IUCrJ. 2020 Jul 1;7(4):728–36. doi:10.1107/s2052252520007411

9. Aumonier S, Engilberge S, Caramello N, von Stetten D, Gotthard G, Leonard GA, et al. Slow protein dynamics probed by time-resolved oscillation crystallography at room temperature. IUCrJ. 2022 Nov 1;9(6):756–67. doi:10.1107/s2052252522009150

10. Kupitz C, Basu S, Grotjohann I, Fromme R, Zatsepin NA, Rendek KN, et al. Serial time-resolved crystallography of photosystem II using a femtosecond X-ray laser. Nature. 2014 Sep 11;513(7517):261–5. doi:10.1038/nature13453

11. Sorigué D, Hadjidemetriou K, Blangy S, Gotthard G, Bonvalet A, Coquelle N, et al. Mechanism and dynamics of fatty acid photodecarboxylase. Science. 2021 Apr 9;372(6538). doi:10.1126/science.abd5687

12. Maestre-Reyna M, Wang PH, Nango E, Hosokawa Y, Saft M, Furrer A, et al. Visualizing the DNA repair process by a photolyase at atomic resolution. Science. 2023 Dec;382(6674). doi:10.1126/science.add7795

13. Rios-Santacruz R, Poddar H, Pounot K, Heyes DJ, Coquelle N, Mackintosh MJ, et al. Integrated structural dynamics uncover a new B12 photoreceptor activation mode. Nature. 2026 Feb 26;650(8103):1045–52. doi:10.1038/s41586-025-10074-2

14. Schulz EC, Yorke BA, Pearson AR, Mehrabi P. Best practices for time-resolved serial synchrotron crystallography. Acta Crystallogr Sect Struct Biol. 2022 Jan 1;78(1):14–29. doi:10.1107/S2059798321011621

15. Monteiro DCF, Amoah E, Rogers C, Pearson AR. Using photocaging for fast time-resolved structural biology studies. Acta Crystallogr Sect Struct Biol. 2021 Oct 1;77(10):1218–32. doi:10.1107/s2059798321008809

16. Schmidt M. Reaction Initiation in Enzyme Crystals by Diffusion of Substrate. Crystals. 2020 Feb 13;10(2):116. doi:10.3390/cryst10020116

17. Schmidt M. Mix and Inject: Reaction Initiation by Diffusion for Time-Resolved Macromolecular Crystallography. Adv Condens Matter Phys. 2013;2013:1–10. doi:10.1155/2013/167276

18. Olmos JL, Pandey S, Martin-Garcia JM, Calvey G, Katz A, Knoska J, et al. Enzyme intermediates captured “on the fly” by mix-and-inject serial crystallography. BMC Biol. 2018 Dec;16(1):59. doi:10.1186/s12915-018-0524-5

19. Calvey GD, Katz AM, Pollack L. Microfluidic Mixing Injector Holder Enables Routine Structural Enzymology Measurements with Mix-and-Inject Serial Crystallography Using X-ray Free Electron Lasers. Anal Chem. 2019 Jun 4;91(11):7139–44. doi:10.1021/acs.analchem.9b00311

20. Pandey S, Calvey G, Katz AM, Malla TN, Koua FHM, Martin-Garcia JM, et al. Observation of substrate diffusion and ligand binding in enzyme crystals using high-repetition-rate mix-and-inject serial crystallography. IUCrJ. 2021 Nov 1;8(6):878–95. doi:10.1107/S2052252521008125

21. Zielinski KA, Dolamore C, Dalton KM, Smith N, Termini J, Henning R, et al. Resolving DJ-1 Glyoxalase Catalysis Using Mix-and-Inject Serial Crystallography at a Synchrotron [Internet]. Biochemistry; 2024 [cited 2026 Aug 4]. Available from: http://biorxiv.org/lookup/doi/10.1101/2024.07.19.604369 doi:10.1101/2024.07.19.604369

22. Stubbs J, Hornsey T, Hanrahan N, Esteban LB, Bolton R, Malý M, et al. Droplet microfluidics for time-resolved serial crystallography. IUCrJ. 2024 Mar 1;11(2):237–48. doi:10.1107/S2052252524001799

23. Mehrabi P, Schulz EC, Agthe M, Horrell S, Bourenkov G, von Stetten D, et al. Liquid application method for time-resolved analyses by serial synchrotron crystallography. Nat Methods. 2019 Oct;16(10):979–82. doi:10.1038/s41592-019-0553-1

24. Butryn A, Simon PS, Aller P, Hinchliffe P, Massad RN, Leen G, et al. An on-demand, drop-on-drop method for studying enzyme catalysis by serial crystallography. Nat Commun. 2021 Jul 22;12(1):4461. doi:10.1038/s41467-021-24757-7

25. Beyerlein KR, Dierksmeyer D, Mariani V, Kuhn M, Sarrou I, Ottaviano A, et al. Mix-and-diffuse serial synchrotron crystallography. IUCrJ. 2017 Nov 1;4(6):769–77. doi:10.1107/S2052252517013124

26. Zielinski KA, Prester A, Andaleeb H, Bui S, Yefanov O, Catapano L, et al. Rapid and efficient room-temperature serial synchrotron crystallography using the CFEL TapeDrive. IUCrJ. 2022 Nov 1;9(6):778–91. doi:10.1107/s2052252522010193

27. 27. Moffat K. Time-Resolved Crystallography. Acta Crystallogr A. 1998 Nov 1;54(6):833–41. doi:10.1107/S0108767398010605

28. Garman EF. Radiation damage in macromolecular crystallography: what is it and why should we care? Acta Crystallogr D Biol Crystallogr. 2010 Apr 1;66(4):339–51. doi:10.1107/S0907444910008656

29. Rajendran C, Dworkowski FSN, Wang M, Schulze-Briese C. Radiation damage in room-temperature data acquisition with the PILATUS 6M pixel detector. J Synchrotron Radiat. 2011 May 1;18(3):318–28. doi:10.1107/S090904951100968X

30. Henzler-Wildman K, Kern D. Dynamic personalities of proteins. Nature. 2007 Dec 13;450(7172):964–72. doi:10.1038/nature06522

31. Indergaard JA, Mahmood K, Gabriel L, Zhong G, Lastovka A, McLeod MJ, et al. Instrumentation and methods for efficient time-resolved X-ray crystallography of biomolecular systems with sub-10 ms time resolution. IUCrJ. 2025 May 1;12(3):372–83. doi:10.1107/S205225252500288X

32. Rolstad AK, Howland E, Sirevåg R. Malate dehydrogenase from the thermophilic green bacterium Chloroflexus aurantiacus: purification, molecular weight, amino acid composition, and partial amino acid sequence. J Bacteriol. 1988 Jul;170(7):2947–53. doi:10.1128/jb.170.7.2947-2953.1988

33. Von Dreele RB. Binding of *N* -acetylglucosamine to chicken egg lysozyme: a powder diffraction study. Acta Crystallogr D Biol Crystallogr. 2001 Dec 1;57(12):1836–42. doi:10.1107/S0907444901015748

34. Beber ME, Gollub MG, Mozaffari D, Shebek KM, Flamholz AI, Milo R, et al. eQuilibrator 3.0: a database solution for thermodynamic constant estimation. Nucleic Acids Res. 2022 Jan 7;50(D1):D603–9. doi:10.1093/nar/gkab1106

35. Synstad B, Emmerhoff O, Sirevåg R. Malate dehydrogenase from the green gliding bacterium Chloroflexus aurantiacus is phylogenetically related to lactic dehydrogenases. Arch Microbiol. 1996 May 22;165(5):346–53. doi:10.1007/s002030050337

36. Talon R, Coquelle N, Madern D, Girard E. An experimental point of view on hydration/solvation in halophilic proteins. Front Microbiol. 2014;5. doi:10.3389/fmicb.2014.00066

37. Becker LM, Fu H, Tatman BP, Dreydoppel M, Kapitonova A, Balazs DM, et al. Aromatic ring flips reveal reshaping of protein dynamics in crystals and complexes. Nat Chem. 2026 Jul;18(7):1221–30. doi:10.1038/s41557-026-02155-0

38. Caramello N, Rose SL, Mathieu E, Petit L, Tews I, Engilberge S, et al. Coupled on-line *in crystallo* UV–Vis absorption spectroscopy and X-ray crystallography to compare specific radiation damage in metal-containing proteins at room versus cryogenic temperature. Acta Crystallogr Sect Struct Biol. 2026 Mar 1;82(3). doi:10.1107/S2059798326000690

39. Bury CS, Brooks-Bartlett JC, Walsh SP, Garman EF. Estimate your dose: RADDOSE-3D. Protein Sci. 2018 Jan;27(1):217–28. doi:10.1002/pro.3302

40. Huang CY, Aumonier S, Engilberge S, Eris D, Smith KML, Leonarski F, et al. Probing ligand binding of endothiapepsin by ‘temperature-resolved’ macromolecular crystallography. Acta Crystallogr Sect Struct Biol. 2022 Aug 1;78(8):964–74. doi:10.1107/S205979832200612X

41. Shimozawa Y, Himiyama T, Nakamura T, Nishiya Y. Structural analysis and reaction mechanism of malate dehydrogenase from *Geobacillus stearothermophilus*. J Biochem (Tokyo). 2021 Sep 22;170(1):97–105. doi:10.1093/jb/mvab027

42. Gotthard G, Aumonier S, De Sanctis D, Leonard G, von Stetten D, Royant A. Specific radiation damage is a lesser concern at room temperature. IUCrJ. 2019 Jul 1;6(4):665–80. doi:10.1107/S205225251900616X

43. Appleby MV, Kepa MW, Winter G, McAuley KE, Beale JH. Modelling of radiation damage and beam-induced heating of room-temperature samples at extremely high flux MX beamlines. IUCrJ. 2026 Mar 1;13(2). doi:10.1107/S2052252525011224

44. Gorel A, Shoeman RL, Hartmann E, Nizinski S, Appleby MV, Beale EV, et al. Testing the limits: serial crystallography using unpatterned fixed targets. IUCrJ. 2025 Nov 1;12(6):692–709. doi:10.1107/S2052252525008371

45. Dalhus B, Saarinen M, Sauer UH, Eklund P, Johansson K, Karlsson A, et al. Structural Basis for Thermophilic Protein Stability: Structures of Thermophilic and Mesophilic Malate Dehydrogenases. J Mol Biol. 2002 May;318(3):707–21. doi:10.1016/S0022-2836(02)00050-5

46. McCarthy AA, Basu S, Bernaudat F, Blakeley MP, Bowler MW, Carpentier P, et al. Current and future perspectives for structural biology at the Grenoble EPN campus: a comprehensive overview. J Synchrotron Radiat. 2025 May 1;32(3):577–94. doi:10.1107/s1600577525002012

47. Wang M. Evolution of macromolecular crystallography beamlines at the Swiss Light Source and SwissFEL. J Synchrotron Radiat. 2025 Sep 1;32(5):1162–83. doi:10.1107/S1600577525005016

48. Kepa MichalW, Tomizaki T, Sato Y, Ozerov D, Sekiguchi H, Yasuda N, et al. Acoustic levitation and rotation of thin films and their application for room temperature protein crystallography. Sci Rep. 2022 Mar 30;12(1). doi:10.1038/s41598-022-09167-z

49. Tsujino S, Sato Y, Jia S, Kepa MW, Trampari S, Tomizaki T. Inertial mixing of acoustically levitated droplets for time-lapse protein crystallography. Droplet. 2024 Jul;3(3):e132. doi:10.1002/dro2.132

50. McGeehan J, Ravelli RBG, Murray JW, Owen RL, Cipriani F, McSweeney S, et al. Colouring cryo-cooled crystals: online microspectrophotometry. J Synchrotron Radiat. 2009 Mar 1;16(2):163–72. doi:10.1107/s0909049509001629

51. von Stetten D, Giraud T, Carpentier P, Sever F, Terrien M, Dobias F, et al. *In crystallo* optical spectroscopy ( *ic* OS) as a complementary tool on the macromolecular crystallography beamlines of the ESRF. Acta Crystallogr D Biol Crystallogr. 2015 Jan 1;71(1):15–26. doi:10.1107/S139900471401517X

52. Caramello N, Adam V, Pearson AR, Royant A. The *in crystallo* optical spectroscopy toolbox. J Appl Crystallogr. 2025 Jun 1;58(3):1068–78. doi:10.1107/s1600576725003541

53. Vonrhein C, Flensburg C, Keller P, Sharff A, Smart O, Paciorek W, et al. Data processing and analysis with the*autoPROC*toolbox. Acta Crystallogr D Biol Crystallogr. 2011 Apr 1;67(4):293–302. doi:10.1107/s0907444911007773

54. Wojdyr M, Keegan R, Winter G, Ashton A. *DIMPLE* - a pipeline for the rapid generation of difference maps from protein crystals with putatively bound ligands. Acta Crystallogr A. 2013 Aug 25;69(a1):s299–s299. doi:10.1107/S0108767313097419

55. Collaborative Computational Project, Number 4. The CCP4 suite: programs for protein crystallography. Acta Crystallogr D Biol Crystallogr. 1994 Sep 1;50(5):760–3. doi:10.1107/s0907444994003112

56. Emsley P, Lohkamp B, Scott WG, Cowtan K. Features and development of *Coot*. Acta Crystallogr D Biol Crystallogr. 2010 Apr 1;66(4):486–501. doi:10.1107/s0907444910007493

57. Liebschner D, Afonine PV, Baker ML, Bunkóczi G, Chen VB, Croll TI, et al. Macromolecular structure determination using X-rays, neutrons and electrons: recent developments in *Phenix*. Acta Crystallogr Sect Struct Biol. 2019 Oct 1;75(10):861–77. doi:10.1107/s2059798319011471

58. Williams CJ, Headd JJ, Moriarty NW, Prisant MG, Videau LL, Deis LN, et al. MolProbity: More and better reference data for improved all-atom structure validation. Protein Sci. 2018 Jan;27(1):293–315. doi:10.1002/pro.3330

